# CNVeil resolves haplotype-specific copy number and uncovers subclonal architecture hidden from total copy number profiling in single-cell cancer genomes

**DOI:** 10.64898/2026.07.20.739667

**Authors:** Weiman Yuan, Can Luo, Yunfei Hu, Liting Zhang, Zi-Hang Wen, Yichen Henry Liu, Xian Mallory, Xin Maizie Zhou

**Author notes:** Equal contributor (Weiman Yuan, Can Luo).

## Abstract

Single-cell DNA sequencing (scDNA-seq) resolves copy number variation (CNV) at single-cell resolution, revealing tumor heterogeneity and subclonal structure. Most existing methods, however, infer only total copy number. Haplotype-resolved copy number, which captures allelic imbalance and clonal evolution, remains far less developed, largely because low coverage, allelic dropout, and technical noise in scDNA-seq make phased allelic inference substantially harder than total copy number estimation. We present CNVeil, a haplotype-aware framework that infers total, allele-specific, and chromosome-scale haplotype-resolved copy number from scDNA-seq data. CNVeil first builds robust total copy number profiles through highly variable bin selection, hierarchical clustering, subclone-aware ploidy estimation, and cross-cell consensus segmentation. Using this profile as a stable scaffold, it infers allele-specific copy number with an expectation-maximization algorithm applied to heterozygous SNP allele counts, then reconstructs haplotype-specific copy number by enforcing coherent haplotype orientation across adjacent segments via dynamic programming. We benchmarked CNVeil against 12 state-of-the-art methods, including eight total copy number callers, two allele-specific callers, and two haplotype-resolved callers, across 20 simulated and real datasets spanning six experimental settings, including high-multiplexed single-nucleus sequencing, Acoustic Cell Tagmentation (ACT), and 10x Chromium. This constitutes the largest comparative evaluation of single-cell copy number inference methods to date. CNVeil consistently outperformed existing tools in segmentation accuracy, ploidy inference, subclone identification, and allele-specific copy number estimation. In a breast cancer multi-omics (wellDR-seq) cohort, CNVeil uncovered haplotype-specific subclonal diversification invisible to total copy number analysis alone and linked allele-specific copy number states to transcriptional variation. By transforming sparse single-cell allelic signals into chromosome-scale haplotype-resolved profiles, CNVeil closes a major methodological gap and provides a scalable framework for studying tumor evolution and functional genomic heterogeneity at single-cell resolution.

## Introduction

Copy number variations (CNVs) are large-scale genomic alterations that change the number of copies of DNA segments and represent a major source of genetic variability in human disease. CNVs can occur at multiple scales, ranging from focal amplifications or deletions to chromosome-arm-level events and whole-chromosome aneuploidy, and are frequently accompanied by loss of heterozygosity (LOH). These alterations can perturb gene dosage, disrupt regulatory balance, and bias allelic representation, thereby influencing transcriptional programs and other downstream cellular processes. Tumor cells provide a particularly relevant biological context for studying CNVs, as they harbor abundant and diverse copy number alterations arising from genomic instability during cancer evolution. The resulting complex CNV landscapes contribute to tumor heterogeneity and complicate therapeutic intervention. Because tumors represent an abundant and dynamically evolving source of copy number variations, understanding how CNVs accumulate during tumor growth requires approaches that capture their evolutionary history. In this context, haplotype-resolved CNVs provide additional information beyond total copy number by retaining allele-level patterns shaped by successive genomic events.

Single-cell DNA sequencing (scDNA-seq) enables genome-wide characterization of copy number alterations at single-cell resolution [1, 2], providing a detailed view of tumor clonal architecture. By avoiding the averaging effects inherent to bulk sequencing, scDNA-seq facilitates the detection of CNV heterogeneity across tumor subclones, supports subclone identification through integrative analyses of genomic alterations such as SCGclust [3], and enables reconstruction of tumor evolutionary trajectories using approaches ranging from copy-number phylogenies to longitudinal evolutionary models [4–9] . More broadly, complementary single-cell modalities have also advanced the characterization of intratumoral heterogeneity. While scDNA-seq directly profiles genomic alterations, scRNA-seq-based approaches such as DeepMalignant identify malignant cell populations from transcriptional programs, providing an orthogonal perspective on tumor heterogeneity [10].In contrast to single-cell RNA sequencing, which is restricted to transcribed regions, scDNA-seq interrogates the entire genome and thus offers a more direct representation of copy number alterations [11–14]. However, scDNA-seq data are susceptible to technical biases, including low and non-uniform coverage, amplification artifacts, and allelic dropout, which together hinder accurate CNV detection and interpretation [15, 16].

A variety of computational methods have been developed to infer total CNVs from scDNA-seq data. Early approaches such as HMMcopy [17], Ginkgo [18], and AneuFinder [19] primarily focus on total copy number inference using Hidden Markov Models or circular binary segmentation. More recent multi-cell methods, including SCOPE [12], SCCNAInfer [20], SeCNV [21], SPRINTER [22], FLCNA [23], and deep learningbased approaches such as rcCAE [24], leverage shared information across cells to improve robustness against technical noise. Although these methods demonstrate improved performance for total copy number estimation, they generally do not resolve allele-specific or haplotype-specific copy number alterations.

Total copy number profiles alone may obscure biologically important events such as loss of heterozygosity at tumor suppressor loci or haplotype-restricted oncogene amplifications, which can drive subclonal diversification and therapy resistance [25, 26]. Resolving haplotype-specific CNVs, distinguishing copy number changes between homologous chromosomes, therefore provides additional insight into allele-level selection and tumor evolution. However, accurate haplotype-resolved CNV inference from scDNA-seq data remains challenging due to sparse allele coverage, phasing ambiguity, and the sensitivity of allele-frequency signals to technical noise [11].

Several methods have been developed to infer allelespecific or haplotype-specific CNV inference from scDNA-seq data, including CHISEL [27], Alleloscope [28], SEACON [29], and CNRein [30]. These methods represent important advances toward resolving allelic copy number variation at single-cell resolution. However, accurate inference remains challenging because scDNA-seq data are characterized by low coverage, sparse SNP observations, and substantial allelic dropout. In particular, CHISEL and Alleloscope jointly infer total and allele-specific copy number states using both read-depth ratios and sparse B-allele frequency (BAF) signals derived from heterozygous SNPs. Consequently, noise in either signal can propagate through the coupled inference procedure, potentially compromising the accuracy of both total and allele-specific copy number estimates.

In addition, CHISEL and CNRein rely on external reference panels or database-based phasing approaches to infer haplotype structure. Because such approaches may not fully capture sample-specific haplotype configurations, phasing inaccuracies can propagate to downstream allele-specific copy number inference. Alleloscope and SEACON infer allele-specific copy number but do not reconstruct chromosome-scale haplotype-specific copy number profiles, preventing the integration of allelic states across genomic segments into coherent haplotypes. Furthermore, with the exception of CNRein, existing methods generally do not explicitly enforce genome-wide consistency of copy number alteration events across chromosomes. As a result, inferred copy number profiles may contain fragmented or biologically implausible alteration patterns, which can reduce the robustness of subclone identification and hinder accurate reconstruction of tumor evolutionary histories.

To address these limitations, we developed CNVeil, a haplotype-aware framework for copy number inference from scDNA-seq data. CNVeil infers total copy number and allele-specific copy number through a hierarchical two-stage procedure. It first leverages the more robust read depth signal to infer total copy number, which then serves as a constraint for allelespecific copy number inference. This formulation substantially reduces the search space because both major and minor allele copy numbers are nonnegative integers whose sum must equal the inferred total copy number. For example, if the total copy number of a segment is inferred to be 3, only two allelespecific states are possible: (3,0) and (2,1). After determining allele-specific copy number states for individual segments, CNVeil reconstructs chromosome-scale haplotype-specific copy number profiles under a biologically motivated minimum-event model that enforces coherent haplotype orientation across adjacent segments. This strategy resolves the correspondence between major and minor alleles genome-wide, thereby transforming sparse and noisy allelic signals into coherent haplotype-resolved CNV profiles and revealing latent subclonal genomic structure.

We evaluated CNVeil across both simulated datasets with known ground truth and diverse real scDNA-seq cohorts, including breast cancer T10, T16, KTN302, and large-scale ACT tumors, as well as 10x Chromium data and wellDR-seq multiomics datasets. We benchmarked CNVeil against 12 existing methods spanning total, allele-specific, and haplotype-resolved copy number inference, including eight total copy number callers, two allele-specific callers, and two haplotype-resolved callers, representing the most comprehensive comparative evaluation of single-cell copy number inference methods to date. These analyses systematically assessed CNVeil’s performance in total, allele-specific, and haplotype-resolved copy number inference. Across all datasets, CNVeil consistently achieved accurate and robust copy number profiling while scaling efficiently to large single-cell cohorts. Beyond benchmarking accuracy, we demonstrated that haplotype-resolved copy number provided biological insight that are inaccessible from total copy number alone, revealing subclone-specific allelic diversification, shared ancestral genomic alterations, lineage-specific dosage remodeling, and their associated transcriptional consequences. Together, these results establish CNVeil as a robust and scalable framework for haplotype-resolved copy number inference and a practical tool for dissecting allele-specific tumor evolution from single-cell DNA sequencing data.

## Results

### Overview of CNVeil

CNVeil infers haplotype-resolved copy number through a hierarchical framework composed of four major modules (**Figure 1**): preprocessing, total copy number inference, allele-specific copy number inference, and haplotype-specific copy number reconstruction. In preprocessing, CNVeil extracts GC-corrected bin-level read depth profiles from single-cell BAM files while filtering unstable genomic regions. Using this corrected read-depth matrix, CNVeil applies agglomerative clustering to identify normal and tumor populations and resolve tumor subclones. Based on the hierachical clustering results, CNVeil subsequently estimates subclone-specific ploidy, and performs cross-cell consensus segmentation for robust total copy number inference. Next, CNVeil integrates SNP-level allelic imbalance signals with the inferred total copy number scaffold. An expectation-maximization algorithm is applied to estimate allele-specific copy number states. Finally, CNVeil reconstructs chromosome-scale haplotype-specific copy number profiles by linking allelic states across adjacent genomic segments under a parsimonious minimum-event model. Together, these modules transform sparse and noisy single-cell sequencing signals into coherent total, allele-specific, and haplotype-resolved copy number profiles.

**Figure 1.**
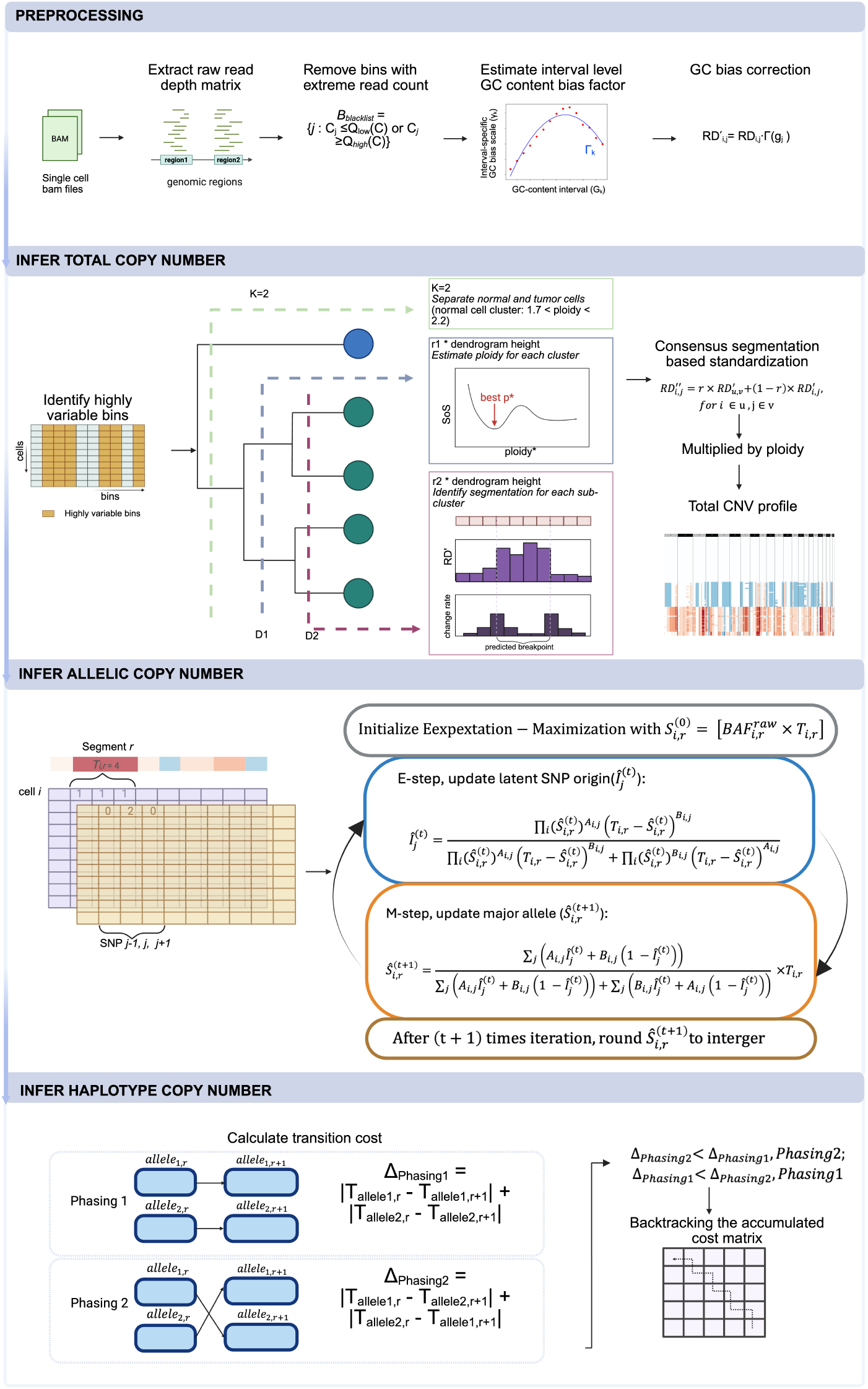
CNVeil workflow. The CNVeil workflow consists of four conceptual modules: 1) *Read depth extraction and GC correction*, which converts aligned single-cell DNA sequencing BAM files into a GC-corrected cell-by-bin read depth matrix; 2) *Total copy number inference*, which uses hierarchical clustering, subclone-aware ploidy estimation, and cross-cell consensus segmentation to infer robust total copy number profiles; 3) *Allele-specific copy number inference*, which leverages SNP-level allele counts within segments and constrains allele-specific states by the inferred total copy number; and 4) *Haplotype-specific copy number reconstruction*, which resolves the orientation of allele-specific states across adjacent segments to generate chromosome-scale haplotype-resolved copy number profiles. Detailed descriptions of each module are provided in the Methods.

### CNVeil robustly recovered copy number profiles across simulated data of various ploidy conditions

We first used the simulated data to evaluate the performance of CNVeil and existing tools (**Table 1**) due to the lack of ground truth data in most real datasets. To simulate datasets and the ground truth, we employed SimSCSnTree [31], which stochastically simulates a tree and imputes CNAs on the tree. Specifically, SimSCSnTree allows users to decide multiple factors such as ploidy of the cells, tree structure, number of clones, number of cells, size of the CNAs, ratio between amplification and deletion, whether there is whole genome duplication, and so on. For the experimental design of our simulations, we simulated four datasets whose average ploidy varying from 1.5 to 5. The number of tumor cells in each dataset was 96, 97, 95, and 100, respectively (**Table 2**). In addition, we also simulated 50 normal cells as a negative control, intermixed with tumor cells as input for all CNV profiling tools. More simulation details are outlined in the Methods section.

**Table 1.** CNV inference methods used in this paper. Sorted by publication year, with implementation language, the finest copy number level resolved (total/allele/haplotype), version, reference, and code link.

| Method | Language | CN level | Version | Reference | Code |
| --- | --- | --- | --- | --- | --- |
| HMMcopy | R | Total | 1.40.0 | Shah et al. 2006 | https://github.com/shahcompbio/HMMcopy |
| Ginkgo | R | Total | default | Garvin et al. 2015 | https://github.com/robertaboukhalil/ginkgo |
| AneuFinder | R | Total | 1.27.1 | Bakker et al. 2016 | https://github.com/ataudt/aneufinder |
| SCOPE | R | Total | 1.10.0 | Wang et al. 2019 | https://github.com/rujinwang/SCOPE |
| CHISEL | Python | Haplotype | 1.0.0 | Zaccaria et al. 2021 | https://github.com/raphael-group/chisel |
| Alleloscope | R | Allele | 1.0.1 | Wu et al. 2021 | https://github.com/seasoncloud/Alleloscope |
| SeCNV | Python | Total | default | Wang et al. 2022 | https://github.com/deepomicslab/SeCNV |
| rcCAE | Python | Total | default | Yu et al. 2023 | https://github.com/zhyu-lab/rccae |
| FLCNA | R | Total | 1.0 | Fei et al. 2024 | https://github.com/FeifeiXiao-lab/FLCNA |
| SEACON | Python | Allele | 1.1.0 | Weiner et al. 2024 | https://github.com/NabaviLab/SEACON |
| SPRINTER | Python | Total | 1.0.0 | Lucas et al. 2024 | https://github.com/zaccaria-lab/SPRINTER |
| CNRein | Python | Haplotype | default | Ivanovic et al. 2025 | https://github.com/elkebir-group/CNRein |
| CNVeil | Python | Haplotype | default | This study | https://github.com/maiziezhoulab/CNVeil |

**Table 2.**
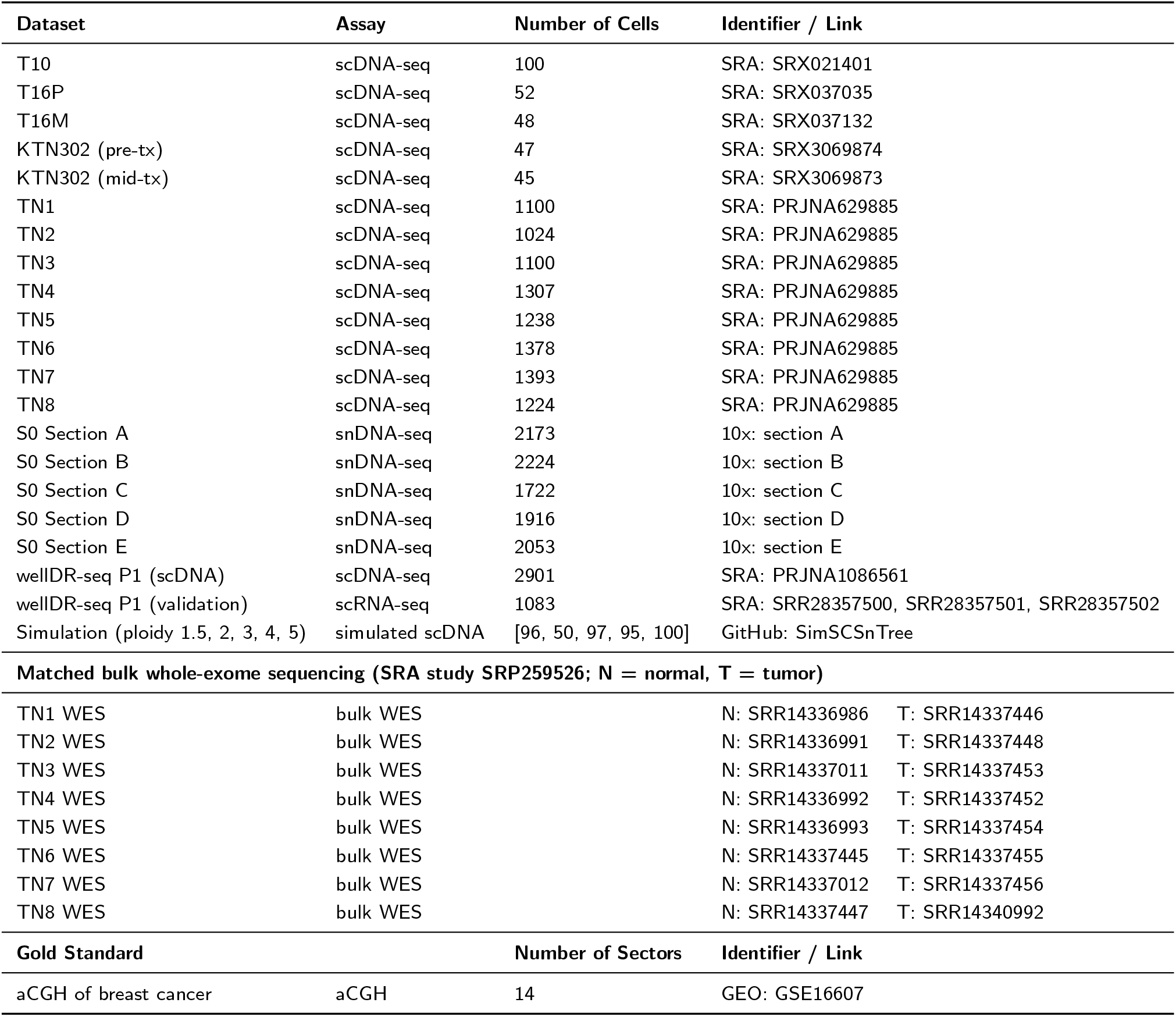
Datasets and gold standard used in this paper. Real and simulated scDNA-seq datasets (with scRNA validation and matched bulk WES where available), giving assay type, number of cells, and identifier/link; for simulated data the simulator is listed. The bottom panel gives the aCGH gold standard.

When assessing the CNV profile generated by each tool against the gold standard, we employed two different evaluation modes following the practice of [32]: segmentation mode and CNV mode. In segmentation mode, our comparison focused solely on the position of breakpoints (segmentation boundaries). If a tool accurately identified the breakpoint where the copy number changed, it was considered a correct breakpoint. In stringent CNV mode, an inference was regarded as correct if the tool identified both the correct breakpoint and the correct copy number for the left and right bins of the breakpoint. The evaluation was specifically applied to tumor cells. More details on the evaluation are also outlined in the methods section.

Overall, CNVeil demonstrated superior and robust performance compared with the other eight tools, AneuFinder, Ginkgo, SeCNV, SPRINTER, SCOPE, HMMcopy, FLCNA, and rcCAE across all four simulated datasets spanning different average ploidy levels. CHISEL, CNRein, and SEACON were excluded from this analysis since SNP information was not incorporated into the simulated datasets. Specifically, as shown in the violin plot (**Figure 2a**), CNVeil consistently achieved the highest average recall, precision, and F1 scores across nearly all evaluated conditions, while maintaining a relatively low variation across cells. For the dataset with an average ploidy of 1.5, SCOPE achieved the best recall in the segmentation evaluation, followed closely by CNVeil. Notably, existing tools exhibited substantially greater variability in recall, precision, and F1 scores for CNV evaluation when the average ploidy was low (1.5). In constrast, when the average ploidy was higher (4.0 and 5.0), existing methods showed markedly lower recall, precision, and F1 scores compared with CNVeil (**Supplementary Tables 1-2**). This trend was particularly pronounced in CNV-state evaluation, where accurate copy number assignment was required, than in segmentation evaluation, which only assessed breakpoint detection.

**Figure 2.**
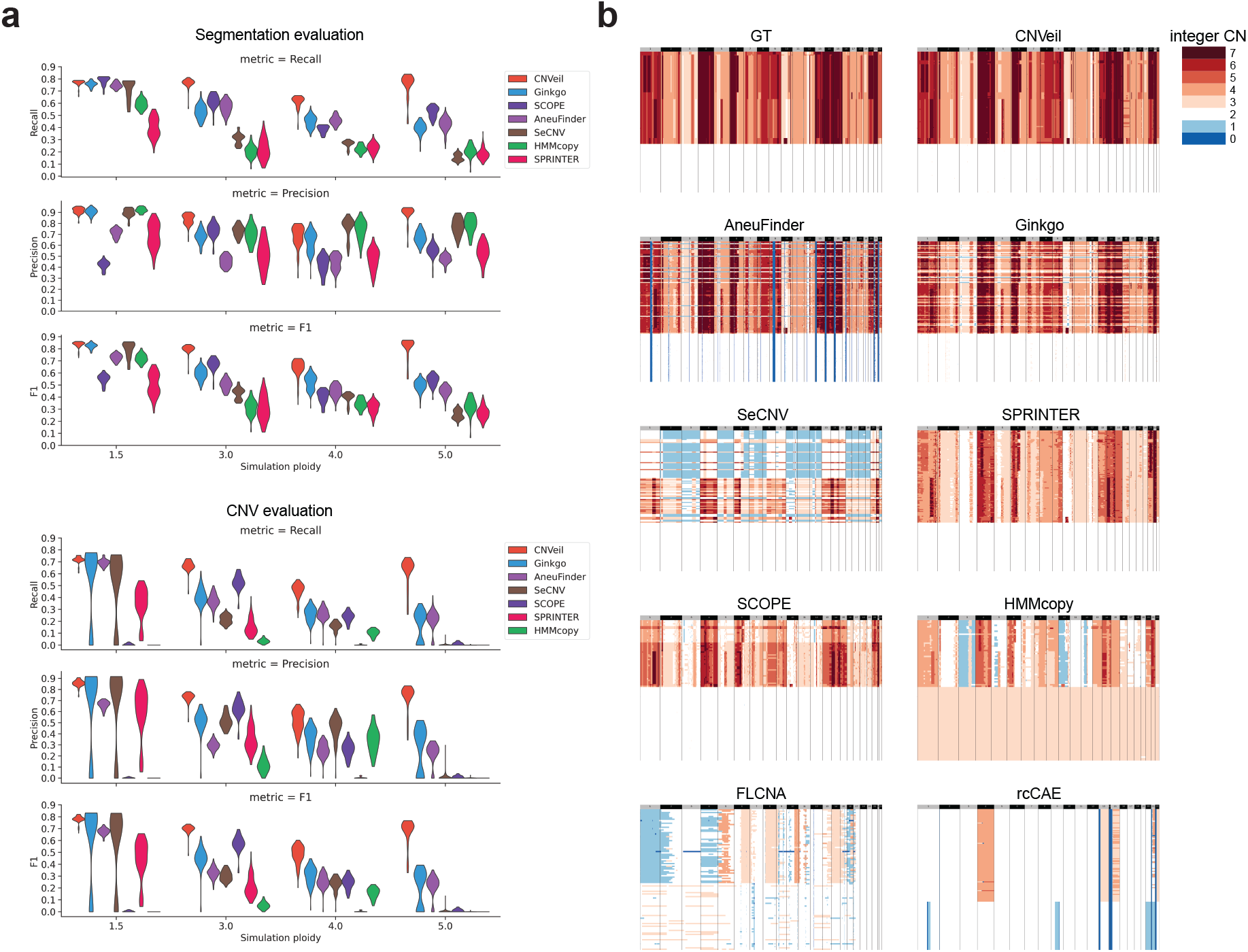
Benchmarking total copy number inference accuracy on simulated single-cell DNA sequencing datasets. (a) Performance comparison of CNVeil and representative single-cell CNV inference methods across simulated datasets spanning multiple ploidy states. Segmentation accuracy (top) and integer copy number inference accuracy (bottom) were evaluated using recall, precision, and F1 score. (b) Representative heatmap comparison of inferred integer copy number profiles for a simulated ploidy = 5 dataset. Ground-truth (GT) copy number states are shown alongside predictions from CNVeil and competing methods. Cells are ordered identically to the ground-truth profile to facilitate direct comparison of chromosomal structure and copy number patterns across methods.

We next visualized the heatmaps of the estimated copy number profiles for all cells across all nine tools in each dataset (**Figure 2b** and **Supplementary Fig. 1**). In all heatmaps, cells were ordered according to the gold-standard phylogenetic tree, such that cells positioned closer together in the heatmap were also more closely related evolutionarily. Consequently, methods that produced more homogeneous copy number patterns across rows were considered to better recapitulate the underlying clonal structure than those that did not. For reference, the heatmap of the groundtruth copy number profile was included as the upperleft panel of each heatmap group.

Across all four simulated datasets, CNVeil produced heatmaps that most closely resembled the groundtruth copy number profiles, demonstrating superior robustness across a wide range of ploidy levels. Specifically, in the high-ploidy dataset (average ploidy = 5.0; **Figure 2b**), CNVeil accurately recovered the overall CNV landscape and clonal structure, while the other eight tools either underestimated tumor-cell ploidy or misassigned copy number states across substantial portions of the genome. In the low-ploidy dataset (average ploidy = 1.5; **Supplementary Fig. 1a**), CNVeil again generated a CNV profile that closely matched the ground truth, while most competing methods systematically overestimated the ploidy of tumor and/or normal cells. For example, SCOPE classified nearly half of the cells as hyperploid in addition to the diploid population, whereas the ground truth contained a hypoploid tumor population together with normal diploid cells. HMMcopy failed to identify the normal-cell population and consistently overestimated tumor-cell copy numbers across the genome. RcCAE largely failed to recover both the tumor clone structure and many underlying copy number alterations. For the intermediate-ploidy datasets (average ploidy = 3.0 and 5.0; **Supplementary Fig. 1b, c**), CNVeil likewise generated CNV profiles that were most consistent with the ground truth. HMMcopy showed the same difficulty in identifying normal cells observed in the low-ploidy dataset. FLCNA underestimated copy numbers across large genomic regions and underestimated the proportion of tumor cells. RcCAE exhibited similar limitations in detecting copy number alterations as observed in the low-ploidy dataset. SPRINTER underestimated copy numbers within tumor subclones in the high-ploidy dataset, whereas SCOPE consistently underestimated the number of tumor cells across all datasets. Taken together, CNVeil was the only method that consistently recovered accurate copy number profiles across all simulated datasets, spanning average ploidy levels from 1.5 to 5.0.

### CNVeil accurately resolved clonal copy number structure in breast cancer patients

We next benchmarked CNVeil against 11 existing tools using real scDNA-seq data obtained from a breast cancer patient identified as T10 [33]. The dataset for patient T10 comprised 100 single cells in total (**Table 2**). Specifically, this dataset incorporated corresponding fluorescence-activated cell sorting (FACS) of the single cell data [34], which revealed four distinct cell subclones: A1 (hyperdiploid, ploidy = 2.85), A2 (hyperdiploid, ploidy = 3.1), H (hypodiploid, ploidy = 1.7), and D (diploid, ploidy = 2). These FACS measurements thus suggested a polygenomic tumor architecture in T10 and provided an independent orthogonal reference for evaluating copy number inference.

The heatmaps in **Fig. 3a** illustrated the estimated total copy numbers across all cells for 12 tools, including those designed for allele-specific copy number inference. To facilitate a direct comparison, cells were ordered according to the FACS-defined populations in all heatmaps. CNVeil, SCOPE, SeCNV, SEACON, AneuFinder, and Ginkgo exhibited broadly similar CNV patterns, identifying a diploid population, a hypodiploid tumor population, and two hyperdiploid tumor populations, consistent with the FACS results.

**Figure 3.**
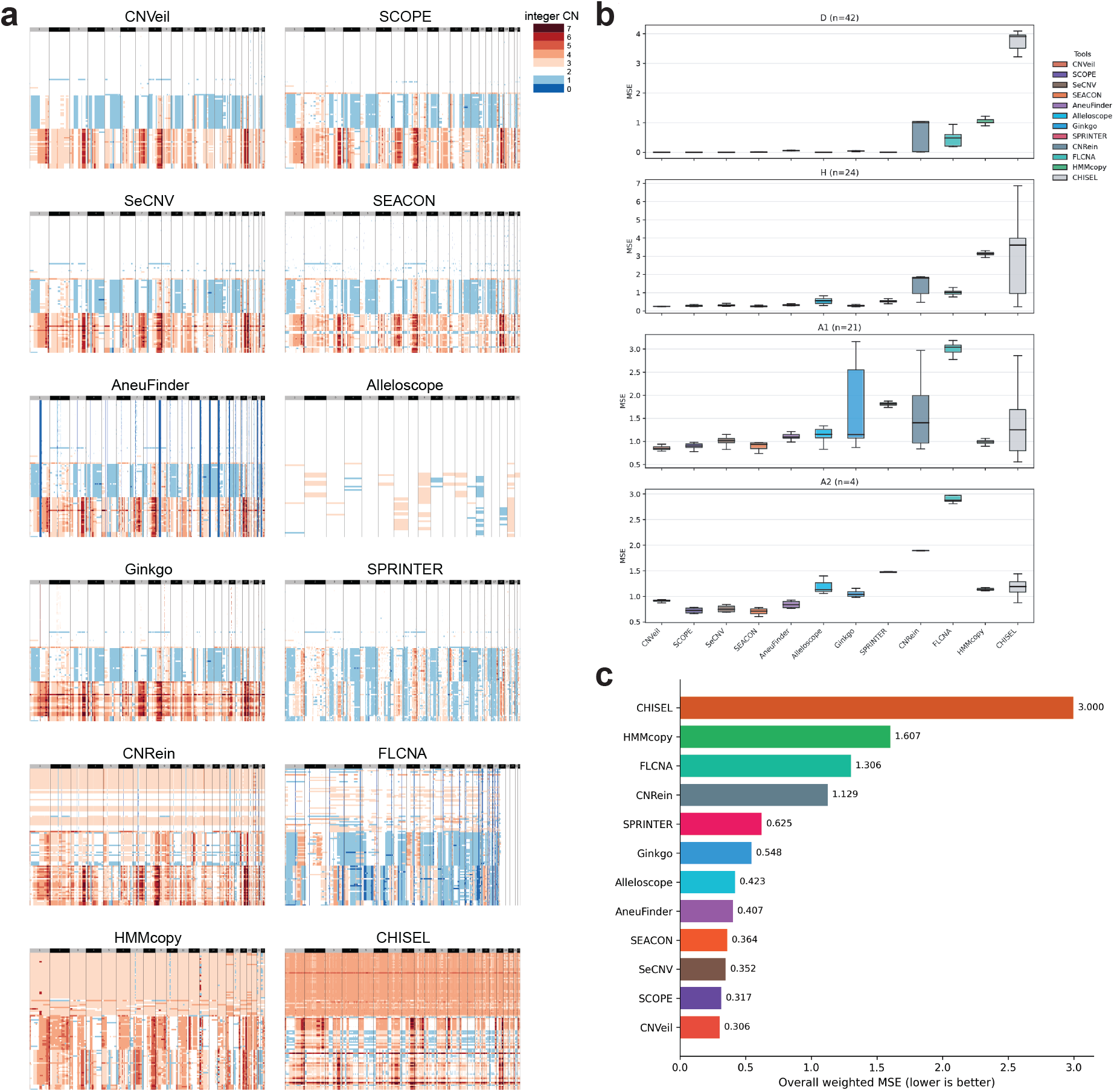
Benchmarking total copy number inference accuracy on the T10 single-cell DNA sequencing dataset. (a) Representative heatmap comparison of inferred total copy number states across methods on the T10 dataset. Columns represent genomic bins ordered by chromosome and rows represent single cells. (b) Mean squared error (MSE) comparison of total copy number inference across four T10 subclones from CNVeil and other callers. Lower MSE indicates improved agreement with ground-truth copy number states. (c) Bar plot showing overall weighted MSE aggregated across T10 four subclones for CNVeil and other callers.

Nevertheless, SeCNV, SEACON, and AneuFinder misclassified CNVeil-defined diploid cells as hypodiploid, with mean genome-wide ploidy estimates of 1.81, 1.82, and 1.84 among the affected cells, respectively. AneuFinder and Ginkgo also exhibited a few outlier bins, suggesting incomplete correction of technical artifacts during preprocessing or downstream inference. Among these tools, CNVeil generated the clearest and most accurate CNV profile for T10. Notably, the two hyperdiploid populations displayed highly similar copy number landscapes, supporting previous observations that the relapse originated from a subclone already present in the primary tumor. CNRein and HMMcopy exhibited similar patterns but systematically overestimated the ploidy of the diploid and hypodiploid subclones. In constrast, FLCNA and SPRINTER tended to underestimate the ploidy of the hyperdiploid subclone. Finally, the allele-specific copy number inference methods Alleloscope and CHISEL produced copy number profiles that showed substantially weaker concordance with the FACS-defined populations.

To quantitatively evaluate the performance, we used CNV calls derived from array comparative genomic hybridization (aCGH) of purified bulk samples from the same patient T10 [33] as the gold standard. The aCGH data provided relative copy number information for each bulk sample. For each FACS-identified subclone, a corresponding gold standard CNV profile was established based on the product of the subclone’s ploidy and the relative copy number from aCGH data. We then compared the CNV profiles inferred by each method with these gold standard profiles using mean squared error (MSE) as the evaluation metric (**Supplementary Table 3**). As shown in **Fig. 3b**, CNVeil and SCOPE achieved the lowest MSE in three of the four FACS-defined populations, demonstrating the highest overall concordance with the orthogonal reference data.

To summarize performance across the entire T10 dataset, we computed an overall wrighted MSE for each methods by aggregating across the four sublcones, weighting each subclone’s contribution by its relative cell proportion (**Fig. 3c**). This metric confirmed the treands observed in the per-subclone analysis. CNVeil achieved the lowest overall weighted MSE (0.306), followed by SCOPE (0.317). SeCNV (0.352), SEACON (0.364), AneuFinder (0.407), and Alleloscope (0.423) also achieved low error rates and showed strong agreement with the FACS-defined subclonal structure. Ginkgo (0.548) and SPRINTER (0.625) followed with moderately higher errors. The remaining methods showed substantially lower concordance. CNRein (1.129), FLCNA (1.306), and HMMcopy (1.607) produced larger deviations from the expected copy number profiles, consistent with the ploidy shifts observed in their heatmaps. CHISEL yielded the highest error (3.000), indicating limited agreement between its allele-specific copy number calls and the FACS-defined subclones. Overall, these results demonstrate that CNVeil provides the most accurate and consistent total copy number estimates across the heterogeneous subclonal landscape of patient T10.

To further assess the generalizability of CNVeil, we evaluated two additional scDNA-seq datasets from triple-negative breast cancer patients, KTN302 and T16 (**Supplementary Fig. 2**). Across both datasets, CNVeil consistently recovered chromosome-scale copynumber profiles while preserving the major subclonal structure. In contrast, competing methods exhibited varying degrees of ploidy distortion, oversegmentation, or loss of subclonal resolution, particularly in diploid cell populations. A detailed comparison of individual methods on these datasets is provided in the Supplementary Results section.

### CNVeil maintained accurate ploidy inference in large-scale ACT and 10x Chromium datasets

We further evaluated CNVeil on large-scale single-cell DNA sequencing data from the Acoustic Cell Tagmentation (ACT) cohort, which comprise eight tumor samples with approximately one thousand cells each (**Table 2**). Because these samples contain no normal cells, this cohort also provides an ideal setting to evaluate whether the methods’ performance depends heavily on the presence of diploid cells in the sample.

Performance was evaluated using mean absolute error (MAE), which measures the average deviation from the FACS ploidy estimate, and Pearson correlation coefficient (*r*), which quantifies concordance across samples. Ploidy accuracy was quantified by comparing inferred genome-wide DNA content against fluorescence-activated cell sorting (FACS) measurements obtained independently for each sample. FACS estimates the average DNA content of tumor cells and therefore provides an orthogonal measurement of sample ploidy that is independent of copy number inference algorithms. Because integer copy number states are defined relative to a global ploidy baseline, errors in ploidy estimation affect the whole genome, causing amplified regions to appear copy number neutral or compressing true copy number losses toward lower states.

CNVeil showed the closest agreement with experimentally measured FACS ploidy across all ACT samples (MAE = 0.047, *r* = 0.996; **Fig. 4a**), with every estimate within 0.09 copies of the ground truth. SPRINTER, Ginkgo, SeCNV, and CHISEL generally preserved the overall ploidy trend, but each exhibited sample-specific deviations. SPRINTER achieved good overall concordance (MAE = 0.131, *r* = 0.828), with its largest deviation observed in TN2 (3.58 vs. 3.03). Ginkgo showed similar performance (MAE = 0.155, *r* = 0.733) but underestimated TN4 (3.06 vs. 3.76). SeCNV remained close to the FACS measurements for most samples, but TN4 was substantially underestimated (2.65 vs. 3.76). CHISEL exhibited moderate agreement (MAE = 0.218, *r* = 0.658), with overestimation in TN1 (3.65 vs. 3.45) and underestimation in TN4 (2.97 vs. 3.76) (**Supplementary Fig. 3**). In contrast, the remaining methods exhibited systematic calibration bias rather than isolated outliers. Specifically, AneuFinder, CNRein, HMMcopy, and FLCNA consistently overestimated or underestimated copy numbers across the majority of samples.

**Figure 4.**
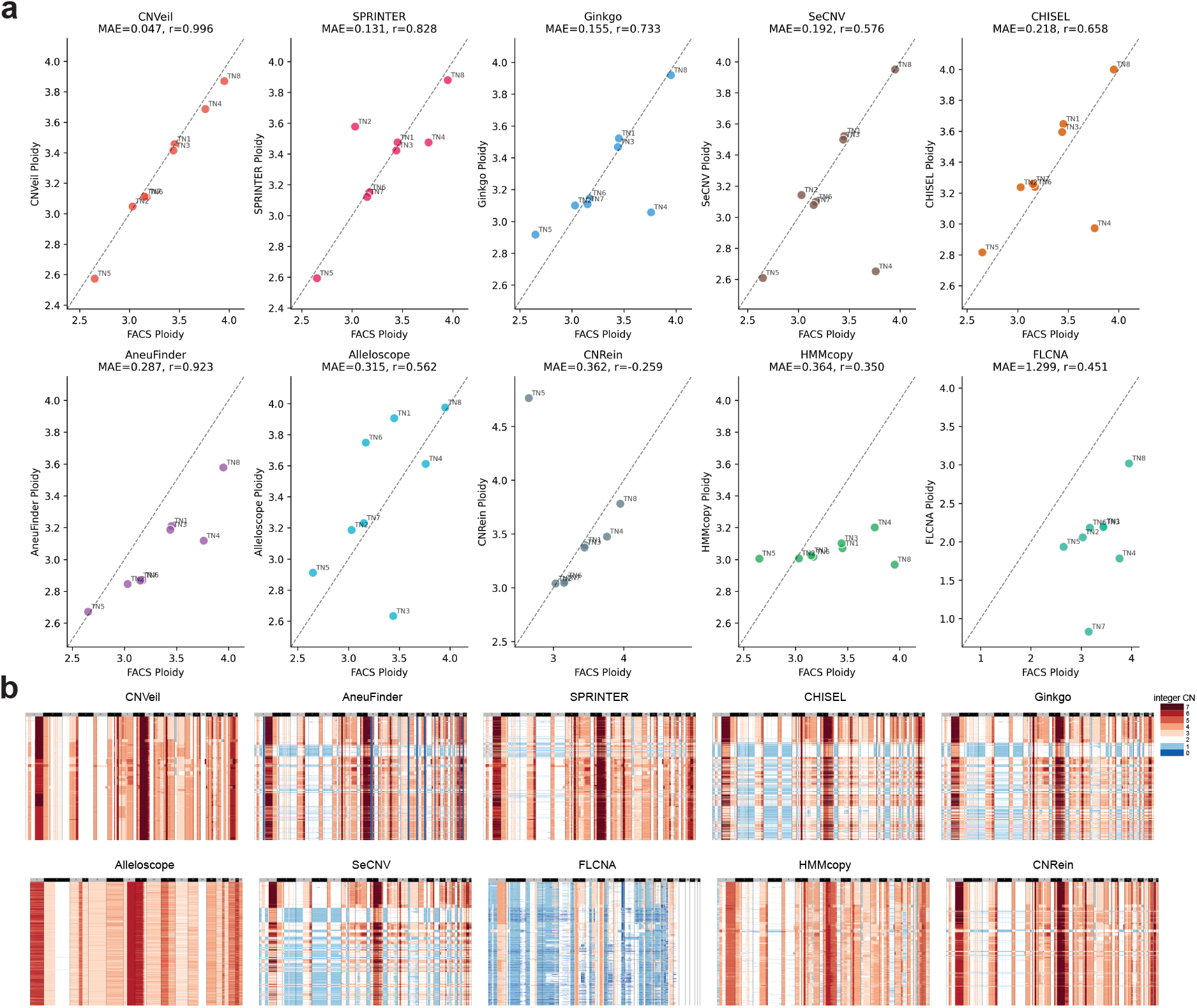
Benchmarking total copy number inference accuracy across ACT single-cell DNA sequencing datasets. (a) Comparison of inferred ploidy estimates against experimentally measured FACS ploidy across ACT tumor samples for CNVeil and representative single-cell CNV inference methods. Each point represents one ACT sample. Mean absolute error (MAE) and Pearson correlation (*r*) relative to FACS measurements are shown for each method. (b) Representative heatmap comparison of inferred integer copy number profiles across ACT tumors. Columns represent genomic bins ordered by chromosome and rows represent single cells. Cells are ordered identically across methods to facilitate direct comparison of chromosomal copy number structure.

The effect of an inaccurate ploidy estimate was directly reflected in the copy number heatmaps for all eight ACT samples (**Fig. 4b** and **Supplementary Fig. 4**). TN4, a hyperploid sample with FACS ploidy of 3.76, was a challenging case in which most tools underestimated the ploidy (**Fig. 4b**). CNVeil accurately recovered genome-wide copy number states of 3-4, yielding a coherent clonal structure without spurious copy number 1 calls. In contrast, SPRINTER, CHISEL, Ginkgo, SeCNV, and CNVRein produced extensive false copy number 1 regions across the genome, while AneuFinder introduced focal copy number 0 segments. FLCNA inferred an overall ploidy of 1.79 for this sample, resulting in copy number 1 assignments across more than half of the genome in most cells.

Similar trends were observed in the remaining ACT samples (TN1-TN3 and TN5-TN8; **Supplementary Fig. 4**), indicating that the observations from TN4 were representative of the cohort rather than sample specific. Overall, CNVeil demonstrated the most robust and accurate copy number profiles across all samples.

We further analyzed a large-scale 10x Chromium dataset S0 comprising five sections (A-E) from patient S0, each containing approximately 2,000 cells (**Supplementary Fig. 5**). Section A consisted predominantly of normal cells, whereas Sections B-E contained progressively larger populations of tumor cells. Among these methods, SPRINTER, CHISEL, and Ginkgo showed the highest agreement with CNVeil, recovering most chromosome scale alterations across the tumor sections. Consistent with its behavior in other datasets, AneuFinder frequently introduced narrow copy number 0 segments throughout the genome, whereas Alleloscope generated broader copy number segments with smoother transitions between adjacent regions. SeCNV tended to overestimate copy number states on chromosome 19 relative to other methods, while SCOPE underestimated copy number states across the proximal region of chromosome 16. Both HMMcopy and CNRein inferred higher copy number states in the diploid region across five sections, suggesting inaccurate normal cell ploidy estimation. Together, these results highlighted the importance of accurate ploidy estimation and demonstrated the robustness of CNVeil across diverse large-scale single-cell DNA sequencing datasets.

**Figure 5.**
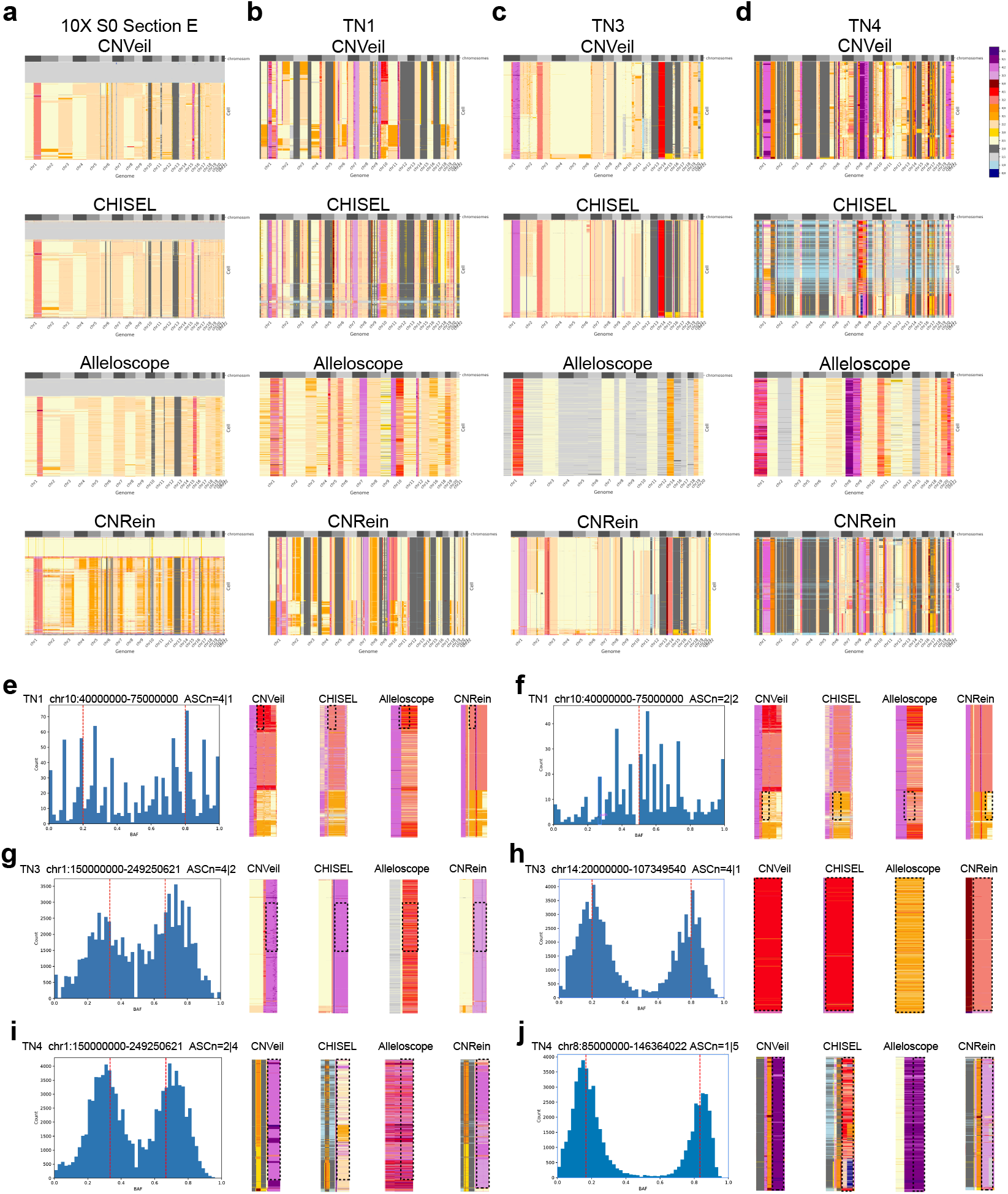
Comparison of allele-specific copy number inference across large-scale single-cell DNA sequencing datasets. (a-d) Representative heatmap comparison of inferred allele-specific copy number states across methods for (a) 10x S0 Section E, (b) ACT TN1, (c) ACT TN3, and (d) ACT TN4. Columns represent genomic bins ordered by chromosome and rows represent single cells. Cells are ordered identically across methods to facilitate direct comparison of allelic structure and subclonal organization. (e-j) Representative genomic regions illustrating allele-specific copy number states supported by B-allele frequency (BAF) distributions. Histograms show BAF distributions for selected regions, with dashed red lines indicating the expected allelic proportions under the inferred allele-specific copy number states. Corresponding heatmaps display inferred allele-specific copy number profiles across methods for the highlighted regions. (e, f) TN1 chr10 (40-75 Mb) containing ASCN states of 4|1 and 2|2. (g, h) TN3 chr1 (150-249.3 Mb) and chr14 (20-107.3 Mb) containing ASCN states of 4|2 and 4|1, respectively. (i, j) TN4 chr8 (85-146.4 Mb) containing an ASCN state of 1|5.

### CNVeil consistently recovered allele-specific copy number states across large-scale datasets

Building on the total copy number evaluation above, we further assessed allele-specific copy number (ASCN) performance on these large-scale datasets. Among the 12 tools benchmarked, only CNVeil, CHISEL, Alleloscope, SEACON, and CNRein were capable of generating ASCN. Because SEACON showed limited concordance with the total copy number patterns recovered by other methods, subsequent ASCN analyses focused on CNVeil, CHISEL, Alleloscope, and CNRein. To facilitate direct comparison, cells were ordered identically across all methods using clustering derived by CNVeil.

In the 10x S0 section E sample, where SNP coverage was relatively balanced, most methods recoverd broadly consistent large-scale clonal structure and ASCN patterns across much of the genome (**Fig. 5a**). Within this overall concordance, however, localized discrepancies became more apparent. CNRein showed systematic deviations, misclassifying diploid regions expected to be (1,1) as elevated copy number states (e.g., total copy number = 3), and introducing asymmetric configurations such as (1,3) in regions where other methods consistently inferred balanced states (e.g., (2,2) in chr4, chr5, and chr12). Alleloscope exhibited a distinct pattern characterized by coarser segmentation and a shifted state; for example, on the first half of chr16, many tumor cells were assigned (3,2), deviating from the (4,2) states inferred by CNVeil and CHISEL. These differences became more apparent in regions with strong allelic imbalance, where small shifts in allele assignment substantially altered the inferred ASCN state.

More pronounced differences were observed in the ACT cohort (**Fig. 5b-d**). A consistent limitation of Alleloscope was apparent across TN samples, where many LOH events were not recovered and regions with strong allelic imbalance were frequently assigned more balanced allele-specific copy number states. In TN1 (**Fig. 5b**), CNVeil, CHISEL, and CNRein recovered broadly similar large-scale structure; however, closer examination at representative loci revealed discrepancies. On chr10 (highlighted using dashed box in **Fig. 5e; heatmaps**), CNVeil identified a segment with allele-specific copy number (4,1), which was well supported by the corresponding BAF histogram showing two peaks centered around 0.2 and 0.8 (**Fig. 5e; histogram**). In contrast, CHISEL and CNRein assigned a more balanced configuration (3,2), while Alleloscope produced a mixture of states across cells, including (3,2), (4,1), and (3,1), indicating reduced stability in allele assignment. For another subclone, in regions corresponding to subclones with balanced SNP counts (**Fig. 5f**), where the BAF distribution showed a single peak near 0.5, CNVeil recovered the expected balanced configuration, whereas CHISEL and CNRein tended to shift toward asymmetric states such as (3,1), and Alleloscope inferred elevated total copy number of 5. A similar pattern was observed on chr1 in TN3 (**Fig. 5g**), where BAF histograms exhibited peaked near 0.33 and 0.67, supporting an imbalanced configuration (4,2) rather than a balanced (3,3). CNVeil and CHISEL captured this state with consistent segment structure, while Alleloscope showed shifted amplitudes, and CNRein more frequently favored balanced interpretations (3, 3). On chr14 (**Fig. 5h**), the BAF distribution supported an ASCN state of (4,1). CNVeil and CHISEL recovered this configuration, whereas Alleloscope substantially reduced the allelic imbalance and CNRein again tended toward more balanced states. In TN4 chr1 (**Fig. 5i**), the BAF distribution exhibited peaked near 0.33 and 0.67, supporting an imbalanced configuration of (2,4). CNVeil consistently preserved this state across most cells. In contrast, CHISEL fragmented the region into mixed states, including (1,2), (1,3), and (2,2), reducing consistency in total copy number across neighboring cells. Alleloscope shifted many cells toward inflated total copy number states, including (2,3), (3,3), and higher-amplitude configurations inconsistent with the observed BAF peaks. CNRein partially recovered the imbalance but frequently alternated between the imbalanced (2,4) state and the balanced (3,3) state, despite clear evidence of asymmetric allele frequency signal. In TN4 chromosome 8 (**Fig. 5j**), the BAF distribution exhibited strongly separated peaks near 0.17 and 0.83, providing clear support for a highly imbalanced state of (1,5). CNVeil consistently preserved this configuration across cells.

The same trends were observed in TN6 and TN8 (**Supplementary Fig. 6**). In TN6, BAF distributions supported multiple levels of allelic imbalance, including states such as (1,3) and (1,4), whereas TN8 contained both highly imbalanced regions and complete LOH events such as (0,2). Across these regions, CHISEL introduced increased local fragmentation, Alleloscope compressed allelic contrast and shifted toward smoother near-balanced states, and CNRein more frequently favored balanced allele configurations despite strong asymmetric BAF evidence. Across these global and local examinations, our observations and analyses indicated that CNVeil more consistently recovered both LOH and graded allelic imbalance, with its inference aligned to the underlying allele frequency signal.

### CNVeil-derived haplotype-resolved copy number profiles demonstrated high concordance with matched bulk WES

To provide an orthogonal assessment of haplotype-resolved copy number inference, we compared CNVeil pseudo bulk profiles against matched bulk whole exome sequencing (WES) data across eight ACT tumors. Bulk allele specific copy number profiles were inferred using ASCAT [35] and phased by aligning bulk allelic imbalance patterns with CNVeil derived pseudo bulk profiles.

ASCAT ploidy estimates were highly concordant with FACS measurements in six of eight ACT samples, with relative errors below 5 percent (**Table 3**). TN4 and TN7 were notable exceptions, showing substantially larger deviations and suggesting inaccurate bulk ploidy inference.

**Table 3.** Comparison of bulk ASCAT ploidy estimates with FACS-derived ploidy measurements across ACT samples. Δ is the absolute difference between FACS and ASCAT ploidy estimates; % error is relative to the FACS measurement.

| Sample | FACS | ASCAT | $\vert \Delta\vert $ | % Error |
| --- | --- | --- | --- | --- |
| TN1 | 3.45 | 3.42 | 0.03 | 0.9 |
| TN2 | 3.03 | 2.99 | 0.04 | 1.3 |
| TN3 | 3.44 | 3.51 | 0.07 | 2.0 |
| TN4 | 3.76 | 4.34 | 0.58 | 15.4 |
| TN5 | 2.65 | 2.76 | 0.11 | 4.2 |
| TN6 | 3.17 | 3.05 | 0.12 | 3.8 |
| TN7 | 3.15 | 4.67 | 1.52 | 48.3 |
| TN8 | 3.95 | 3.86 | 0.09 | 2.3 |

We selected TN6 as a representative example because ASCAT and FACS yielded highly concordant ploidy estimates (3.05 and 3.17, respectively). CNVeil pseudo bulk profiles closely matched both total and haplotype specific copy number patterns inferred from bulk WES (**Fig. 6a-b**). Concordance was particularly evident on chromosomes such as chr2, chr10, and chr18, where both approaches consistently recovered the major copy number transitions and haplotype-specific imbalance patterns. Chromosome-level correlations further supported this high degree of agreement (**Fig. 6c**). Correlations were generally highest on chromosomes exhibiting substantial copy number variation, whereas chromosomes with largely flat profiles showed more variable estimates. Reduced, and occasionally negative, correlations were primarily observed on chromosomes with limited copy number variation, where small differences in segment boundaries, local averaging, or haplotype assignment could substantially affect correlation estimates despite broadly concordant copy number profiles.

**Figure 6.**
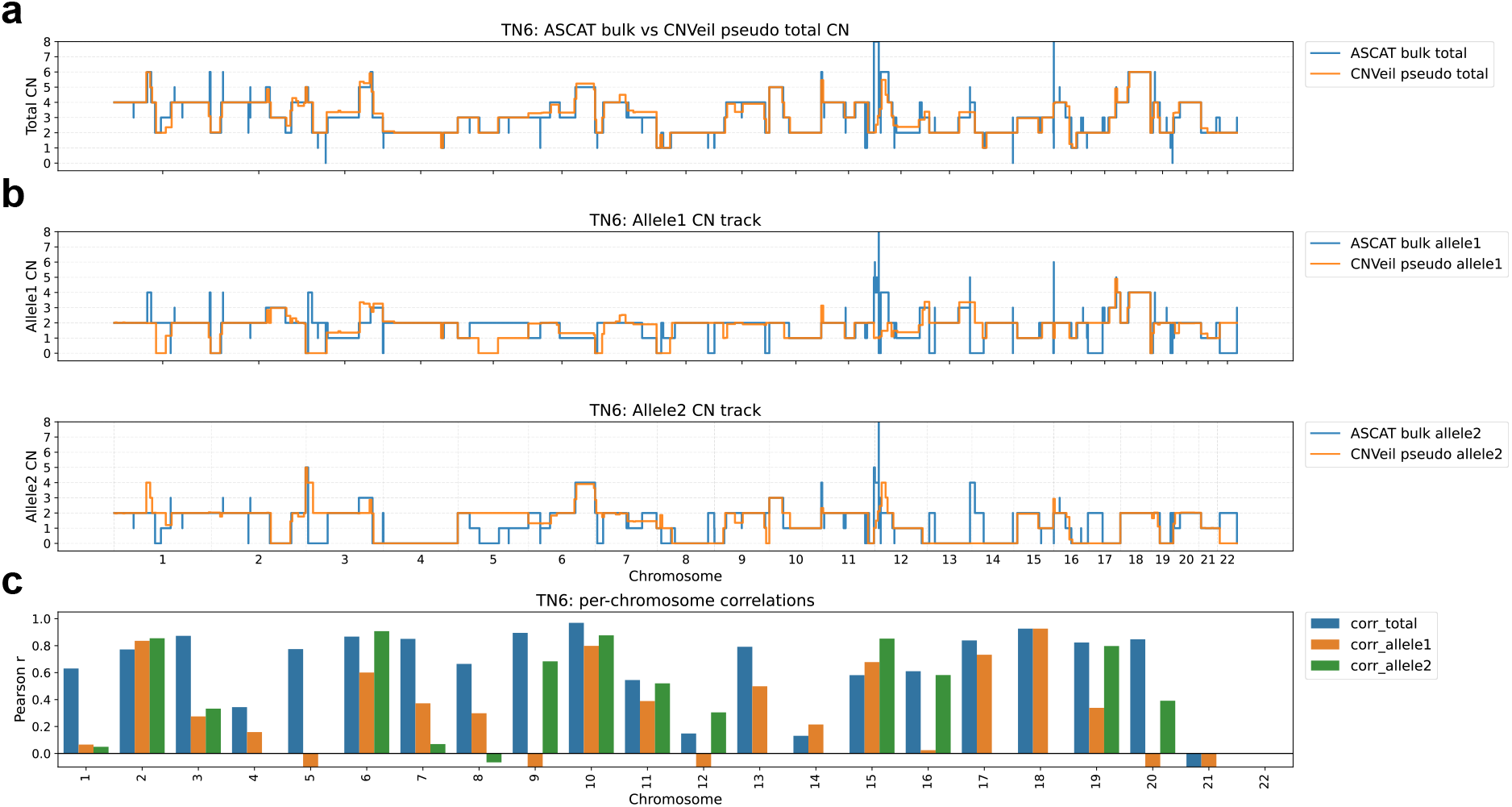
Concordance between CNVeil pseudo-bulk and matched bulk WES haplotype-resolved copy number profiles in ACT sample TN6. (a) Comparison of genome-wide total copy number profiles between phased bulk WES-derived ASCAT estimates and CNVeil pseudo-bulk reconstruction. (b) Comparison of haplotype-specific copy number profiles for the two inferred haplotypes across the genome. (c) Chromosome-level Pearson correlations between pseudo-bulk and bulk-derived profiles for total copy number and haplotype-specific copy number states.

Similar agreement was observed in TN1, TN2, TN3, TN5, and TN8 (**Supplementary Fig. 7a, b, c, e, g**), all of which showed close agreement between ASCAT and FACS ploidy estimates. By contrast, TN4 and particularly TN7 showed weaker agreement across both total and haplotype-specific profiles. Notably, these were also the two samples with the largest discrepancies between ASCAT and FACS ploidy estimates, suggesting that inaccuracies in bulk ploidy inference propagated into downstream allele specific copy number estimates. Together, these analyses provided independent support that CNVeil preserved chromosome-scale haplotype structure inferred from single-cell DNA sequencing data and produced haplotype-resolved copy number profiles broadly consistent with matched bulk WES measurements.

### Haplotype-resolved copy number reveals hidden tumor evolution and lineage-specific genomic programs

To investigate how haplotype-resolved copy number refines tumor evolution beyond total copy number alone, we applied CNVeil to a sample of the wellDR-seq breast cancer cohort and reconstructed phylogenetic trees from consensus copy number profiles of inferred subclones using MEDICC2 [36]. Details for evolutionary and related analysis are provided in the Supplementary Methods section. Consistent with previous single-cell studies, total copy number profiles captured the major genomic architecture of the tumor, reflecting the dominant role of large-scale CNAs in breast cancer evolution [34, 37–39]. However, total copy number obscures allelic configurations that differ in parental origin, LOH status, and haplotype-specific dosage. As a result, phylogenetic reconstruction based solely on total copy number produced a relatively simple topology, whereas reconstruction using haplotype-resolved copy number revealed substantially greater branching complexity (**Fig. 7a**), indicating that distinct evolutionary lineages can remain hidden when tumors are represented only by total copy-number states.

**Figure 7.**
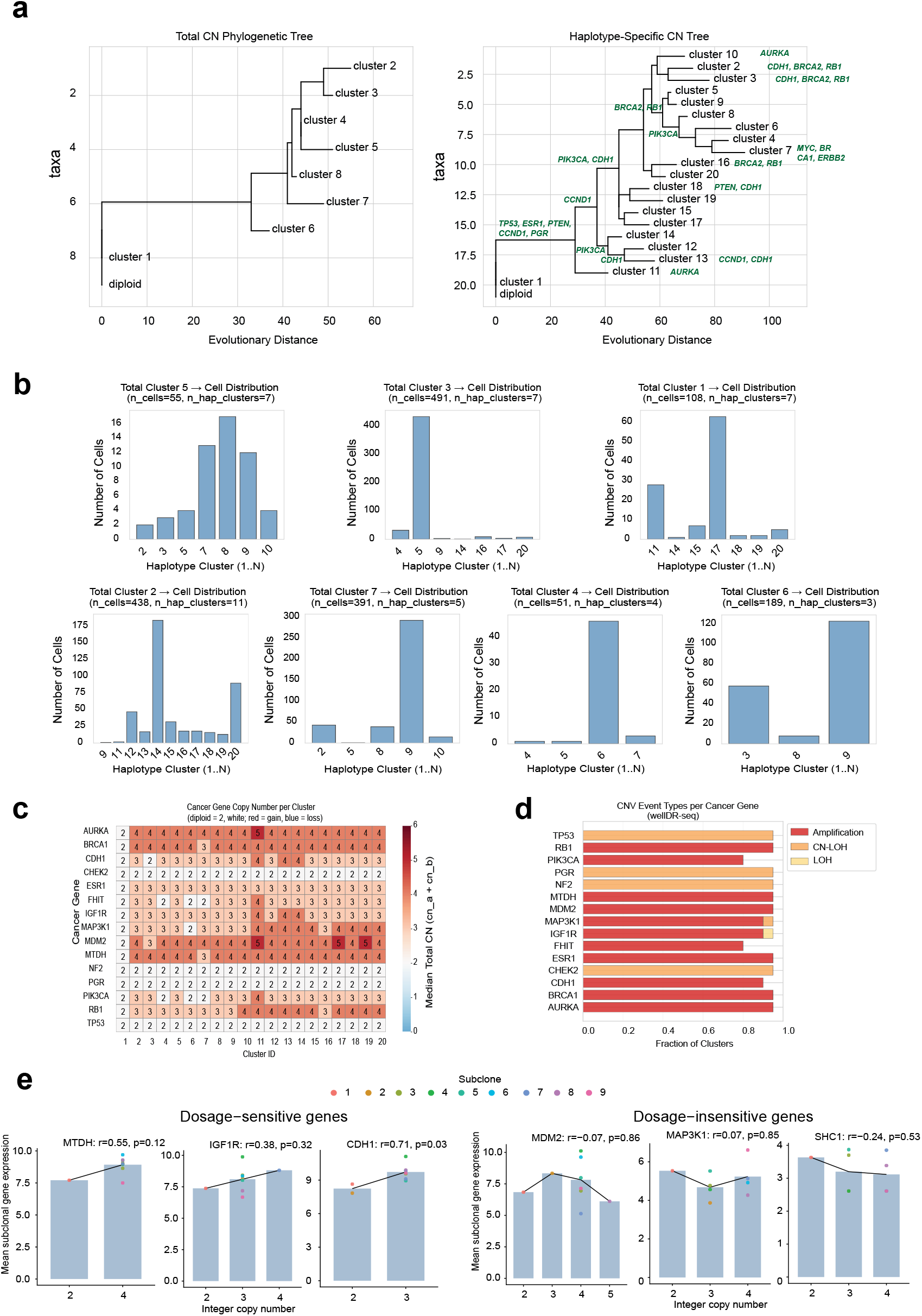
Phylogenetic reconstruction from total and haplotype-resolved copy number profiles in the wellDR-seq breast cancer dataset. (a) Phylogenetic trees reconstructed from total and haplotype-resolved copy number profiles. Representative cancer-associated genes associated with major branches are annotated. (b) Correspondence between total copy number clusters and haplotype-defined clusters. (c) copy number states of representative cancer-associated genes across inferred haplotype-defined subclones. (d) Distribution of amplification, loss of heterozygosity (LOH), and copy-neutral LOH (CN-LOH) events across representative cancer genes. (e) Relationship between inferred copy number dosage and gene expression across inferred subclones for representative genes.

This increased resolution was evident when comparing cluster assignments derived from the two representations (**Fig. 7b**). Individual total copy number clusters frequently subdivided into multiple haplotype-defined groups. For example, total cluster 1, 2, and 3 each split into 7-11 distinct haplotype clusters, whereas total clusters 4 and 6 separated into four and three haplotype groups, respectively. These findings suggest that cells sharing similar total copy number profiles can harbor distinct haplotype architectures, revealing additional evolutionary structure that remains unresolved when tumors are represented solely by total copy number.

This additional evolutionary structure was accompanied by lineage-specific copy number patterns across cancer-associated genes (**Fig. 7c**). Many cancer-associated genes were recurrently altered across multiple inferred lineages, but the corresponding copy number dosage frequently differed between subclones. For example, *PIK3CA, CDH1, RB1*, and *BRCA1* were altered in most lineages while exhibiting distinct copy number states across subclones, illustrating the additional evolutionary resolution revealed by haplotype-resolved copy number profiles.

To further characterize these alterations, we summarized the distribution of amplification, LOH, and copy-neutral LOH events across recurrently altered genes (**Fig. 7d**). Oncogenes such as *AURKA, MDM2, MTDH*, and *PIK3CA* were predominantly affected by amplifications, whereas several tumor suppressor genes more frequently exhibited LOH or copy-neutral LOH. Notably, *TP53* was altered almost exclusively through LOH and copy-neutral LOH, illustrating how biologically important genomic events can occur without changes in total copy number state and therefore remain undetectable by total copy number analysis alone [27, 28, 40, 41].

We next asked whether these lineage-specific copy number configurations were accompanied by corresponding transcriptional changes (**Fig. 7e**). Consistent with previous studies showing that transcriptional responses to copy number alteration are gene dependent rather than universal [42– the relationship between inferred copy number dosage and gene expression varied substantially across genes. For example, *CDH1* exhibited a clear positive association between copy number state and expression across inferred subclones, whereas *MTDH* and *IGF1R* showed similar trends with greater variability. In contrast, *MDM2, MAP3K1*, and *SHC1* displayed comparatively weak correspondence between genomic dosage and transcriptional output, suggesting partial buffering of copy number effects. Together, these results linked haplotype-defined evolutionary lineages to distinct transcriptional programs.

### CPU time and memory analysis

To evaluate the computational efficiency of the benchmarked methods, we recorded the total CPU time and peak memory usage for all available benchmark runs (**Supplementary Table 4**). All tools were executed using single-cell BAM files as input, and the total CPU time was measured from job submission until completion of the final CNV profile. Peak memory usage was obtained from the maximum resident set size (MaxRSS) reported by the Slurm scheduler. Because the benchmark evolved over the course of this study, resource measurements are not available for every tool-dataset combination. Specifically, Ginkgo was initially evaluated through its web interface for the smaller datasets, for which runtime and memory statistics were not accessible, and was therefore profiled only on the larger datasets after the standalone version became available. SCOPE requires a matched normal-cell reference and was therefore not benchmarked on the larger tumor datasets. Conversely, rcCAE was not further evaluated on the larger datasets because of its limited accuracy on the simulated benchmarks. Consequently, Supplementary Table 4 summarizes all available resource measurements rather than a complete tool-by-dataset matrix.

Among the representative large-scale benchmarks, the TN4 dataset (approximately 1,000 cells) illustrates the computational characteristics of the evaluated methods. In terms of computational cost, CNVeil compared favorably against other haplotype- and allele-specific CNV callers. Among the phasing-aware methods, CNVeil completed in 91.87 minutes, substantially faster than CHISEL (534.85 min), SEACON (5400.35 min), and CNRein (82,475.45 min), though slower than the coarser allele-level method Alleloscope (2.03 min). In peak memory usage, CNVeil (98.28 GB) was similar to CHISEL (93.80 GB) and CNRein (91.61 GB), all of which were markedly higher than Alleloscope (4.93 GB). When compared against totalcopy-number callers, CNVeil remained competitive despite performing additional computation; it required more time than HMMcopy (32.26 min), FLCNA (63.88 min), and SeCNV (68.02 min), but was faster than AneuFinder (227.23 min) and SPRINTER (1036.72 min), even though these total-copy-number callers only need to infer total copy number, whereas CNVeil must additionally resolve allele-specific and haplotype-specific copy number on top of the total CNV calling step. This makes CNVeil’s runtime especially favorable given the extra layer of inference it performs.

Unless otherwise specified, all reported resource measurements were obtained using the default benchmark configuration with five CPU threads. Because several stages of CNVeil are parallelized, increasing the number of available threads can substantially reduce wallclock runtime.

## Discussion

Here we present CNVeil, a haplotype-aware framework that infers total, allele-specific, and chromosome-scale haplotype-resolved copy number from single-cell DNA sequencing data. Across simulated datasets spanning average ploidy from 1.5 to 5.0, as well as real cohorts from single-nucleus sequencing, ACT, and 10x Chromium, CNVeil consistently outperformed both total-copy number and allele-aware methods, with its accuracy independently supported by FACS ploidy, aCGH, and matched bulk WES. Existing allele-aware methods such as CHISEL [27] and Alleloscope [28] jointly infer total and allele-specific copy number directly from read-depth ratios and sparse BAF signals, so noise or allelic dropout in either signal propagates into the final states. CNVeil instead establishes a robust total copy number scaffold first and uses it to constrain allele-specific inference within segments of constant dosage, decoupling high-confidence dosage signals from inherently noisy allelic signals. This is most apparent under strong allelic imbalance, where Alleloscope compressed allelic contrast and missed LOH, and CNRein [30] reverted to balanced configurations despite asymmetric BAF, whereas CNVeil recovered LOH and copy-neutral LOH states consistent with the underlying allele frequencies and coherent across neighboring cells. Notably, CNVeil also maintained accurate ploidy inference in cohorts lacking normal cells, demonstrating that its performance does not rely on the presence of a diploid reference population.

Beyond improving copy number inference accuracy, haplotype-resolved analysis provided biological insights that were inaccessible from total copy number alone. In the wellDR-seq breast cancer cohort, total copy number reconstructed a relatively simple evolutionary topology, whereas haplotype-resolved copy number resolved apparently homogeneous clones into multiple evolutionary lineages, revealing allele-specific evolutionary events that remained hidden in total copy number space. These included LOH and copy-neutral LOH affecting tumor suppressor genes such as TP53, events that preserve total copy number while fundamentally altering allelic dosage. Furthermore, integrating haplotype-resolved copy number with matched transcriptional profiles revealed geneand contextdependent dosage effects, highlighting the functional consequences of allele-specific genomic alterations. Together, these results demonstrate that haplotype-resolved copy number provides an additional layer of information for reconstructing tumor evolution and linking genomic alterations to transcriptional heterogeneity.

## Methods

### Design principles

Single-cell DNA sequencing data are characterized by extreme sparsity at the level of individual loci, which makes direct inference of allelic imbalance highly unstable. In particular, B-allele frequency (BAF) measurements at individual SNVs are frequently dominated by sampling noise and allelic dropout, especially in low-coverage regions or genomic segments with limited SNP density. As a result, approaches that attempt to jointly infer total and allele-specific copy number directly from raw BAF and read-depth signals may overfit technical noise and yield biologically implausible copy number configurations.

CNVeil is therefore designed around a hierarchical inference strategy that explicitly separates highconfidence dosage signals from inherently noisy allelic signals. First, total copy number is inferred using aggregated read-depth information across genomic bins and cells, providing a robust estimate of large-scale dosage changes while averaging out locus-level stochasticity.

Second, allele-specific copy number is inferred conditional on the estimated total copy number and constrained within genomic segments of constant dosage. This strategy enables allelic imbalance to be resolved using information from coherent genomic regions rather than relying solely on sparse locus-level BAF measurements. By enforcing consistency between total and allele-specific copy number states, the model substantially reduces the search space and prevents spurious solutions.

Finally, haplotype-specific copy number is inferred globally across genomic segments using a minimum-event criterion. Rather than assigning parental alleles independently at each segment boundary, CNVeil identifies haplotype configurations that minimize the number of copy number alteration events along each chromosome. This parsimony-driven formulation promotes genome-wide coherence and reflects the biological observation that many copy number alterations arise from interval events affecting contiguous chromosomal regions. As a result, inferred haplotypes are not only locally consistent with observed data but also globally consistent with the evolutionary processes that generate large-scale genomic rearrangements.

### CNVeil workflow

CNVeil employs a hierarchical inference framework in which total copy number is first inferred from aggregated read-depth information, followed by allelespecific inference conditioned on total copy number states and global haplotype reconstruction using a minimum-event criterion. The workflow consists of four conceptual stages: 1) read depth extraction and GC correction; 2) total copy number inference; 3) allele-specific copy number inference conditioned on total copy number; 4) haplotype-specific copy number reconstruction via minimum-event optimization.

#### Read depth extraction and GC correction

Read-depth extraction and GC correction in CNVeil are performed using a preprocessing module adapted from AneuFinder [19]. Given aligned single-cell DNA sequencing BAM files as input, CNVeil first partitions the autosomal genome into non-overlapping fixed-width bins, with a default bin size of 500 kb across chromosomes 1-22. For a dataset containing *n* cells and *m* genomic bins, the raw read depth matrix is denoted as

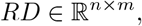

where *RD*_*i*,*j*_ denotes the number of reads from cell *i* assigned to genomic bin *j*. By aggregating reads within genomic bins, CNVeil reduces locus-level sampling variability and provides a more stable readdepth signal for copy number inference in low-coverage single-cell DNA sequencing data.

To remove unstable genomic regions, CNVeil identifies bins with extreme aggregate coverage using a merged BAM file constructed from all cells. Let

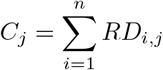

denote the aggregate read count for bin *j*. Bins with aggregate coverage below the lower quantile or above the upper quantile are considered unreliable and excluded from downstream analysis:

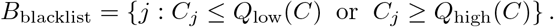

Adjacent blacklisted bins are merged into continuous excluded intervals. This filtering step removes regions exhibiting systematically aberrant coverage patterns, including loci affected by low mappability, repetitive sequences, assembly artifacts, and centromere-proximal mapping errors.

CNVeil then corrects bin-level read counts for GC-content bias. For each genomic bin *j*, let *g*_*j*_ *∈* [0, 1] denote its GC content. Bins are grouped into GC-content intervals *G*_1_, *G*_2_, …, *G*_*K*_, and an interval-specific GC correction scale is estimated as

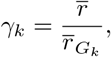

Where 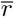 is the genome-wide average read count and 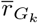 is the trimmed mean read count among bins whose GC content falls within interval *G*_*k*_. Estimating correction factors at the interval level increases robustness by pooling information across bins with similar GC content. A second-order polynomial function is then fitted to the interval-specific GC-bias correction scales, yielding a smooth GC correction curve. The corrected read count for cell *i* and bin *j* is computed as

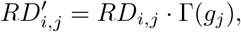

where Γ(*g*_*j*_) denotes the GC correction factor assigned to bin *j* according to its GC content. Corrected counts are rounded to integer values and retained for downstream analysis.

Finally, corrected read depth profiles from all cells are reformatted into a unified cell-by-bin matrix,

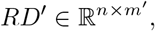

where *m*^*′*^ denotes the number of retained genomic bins after blacklist filtering. Rows correspond to genomic bins labeled by chromosome coordinates, and columns correspond to individual cells. This corrected read depth matrix serves as the input for subsequent processing.

#### Total copy number inference

To infer total copy number, CNVeil uses the GC-corrected read depth matrix as input and converts bin-level read depth signals into integer copy number states through a multi-step procedure consisting of hierarchical clustering, ploidy estimation, consensus breakpoint detection, segmentation, and copy number standardization. First, CNVeil performs hierarchical clustering on highly variable genomic bins to distinguish normal and tumor cells and subsequently identify tumor subclones with shared copy number profiles and ploidy states. For each inferred cluster or subclone, the optimal ploidy is estimated by the value that maximizes the concordance between scaled read-depth measurements and integer copy number states. To reduce cell-level noise and improve breakpoint detection, CNVeil then performs fine-grained clustering within each subclone and identifies consensus breakpoints across cells. Finally, read depth values are standardized using segment-level consensus signals and scaled by the inferred subclone ploidy to generate the final total copy number profile for each cell. More detailed descriptions of these procedures are provided in the following sections.

#### Hierachical clustering for different levels of cell clusters

CNVeil employs agglomerative clustering algorithm [47] at multiple stages of total copy number inference, including normal-tumor cell separation, tumor subclone identification, and fine clustering within subclones. These clustering steps serve two complementary purposes: (i) identifying groups of cells that share similar ploidy states for robust ploidy estimation, and (ii) aggregating information across related cells to improve breakpoint detection and segmentation. Because the absolute copy number of a genomic bin is proportional to the product of normalized read depth and the underlying cellular ploidy, accurate copy number inference requires both reliable ploidy estimation and robust identification of shared copy number breakpoints across cells for subsequent read-depth standardization.

Agglomerative clustering initializes each cell as an individual cluster and iteratively merges clusters according to a similarity measure until all cells are merged into a single cluster. This process generates a tree-like structure, known as a dendrogram, that can be cut at different heights to obtain different clustering solutions at multiple resolutions. Lower cut heights produce finer-grained clusters, whereas higher cut heights produce broader clusters containing more heterogeneous cell populations. The height of a certain level in a dendrogram represents the linkage distance of clusters at that level. CNVeil uses Ward’s linkage criterion to measure the distance between clusters. Ward’s linkage is defined as

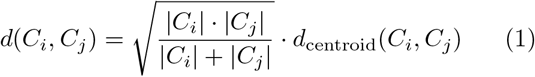

Where d(*C*_*i*_,*C*_*j*_) denotes the Ward’s linkage distance between clusters *C*_*i*_ and *C*_*j*_, *C*_*i*_ and *C*_*j*_ denote the number of clusters contained in the two clusters, and *d*_*centroid*_(*C*_*i*_,*C*_*j*_) denote the Euclidean distance between the centroids of clusters *C*_*i*_ and *C*_*j*_.

#### Initial normal-tumor cell classification

To perform the initial normal-tumor cell clustering, CNVeil uses highly variable genomic bins as features for cell representation rather than the full set of consecutive genomic bins. Highly variable bins exhibit substantial variation in read-depth signals across cells and are therefore more informative for distinguishing distinct cellular populations.

To identify highly variable bins, CNVeil takes the normalized read-depth matrix *RD*^*′*^ as input and computes the variance across cells for each genomic bin. The variance for a given bin *j* can be expressed as:

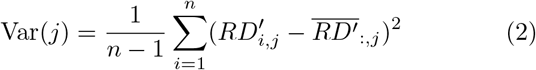

Where 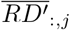 denotes the mean read depth of bin *j* across all cells, *i* is the cell index, and *n* denotes the number of cells. CNVeil selects the top 10% of bins ranked by variance as highly variable bins (HVBs). Because copy number alterations are a major source of read-depth variability, HVBs are enriched for genomic regions that distinguish normal and tumor populations and capture intercellular heterogeneity. The resulting matrix containing only highly variable bins is denoted by 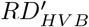, CNVeil then applies agglomerative clustering to 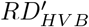, partitioning cells into two clusters by setting the number of clusters to two. Subsequently, CNVeil estimates the optimal ploidy for each cluster using a numerical optimization procedure. The cluster whose estimated ploidy is closest to the diploid state is designated as the normal-cell cluster, whereas the remaining cluster is designated as the tumor-cell cluster.

Specifically, CNVeil identifies the cluster-level ploidy that minimizes the discrepancy between scaled readdepth values and their nearest integer copy number states. This approach is motivated by the expectation that, after appropriate scaling, true copy number states should be close to integer values. The numerical optimization procedure is described below.

CNVeil defines a candidate ploidy set *Ploidy* = [1.50, 1.55, …, 5.50], which spans ploidy values from 1.50 to 5.50 with a step size of 0.05. Let |*P*| = *K*. The Scaled Copy Number Profile (SCNP) for candidate ploidy *k* is defined as:

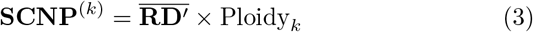

Where 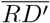 denotes the vector of mean normalized read-depth values across cells for each genomic bin. The corresponding sum of squares deviations (SoS) from the nearest integer copy number states is calculated as:

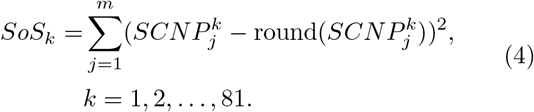

The optimal ploidy level for each cluster, represented by *ploidy*^*\**^ or *Ploidy*[*k*], can be achieved by the value of *k* that minimizes the SoS:

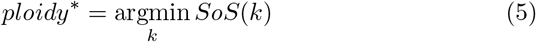

Through this procedure, CNVeil jointly performs normal-tumor cell separation and cluster-level ploidy estimation, providing a robust foundation for downstream breakpoint detection, segmentation, and total copy number inference.

#### Tumor subclones identification and ploidy estimation

In practice, tumor cells from the same patient may comprise multiple subclones that either share a common ploidy state or exhibit distinct ploidy levels [35, 39, 48]. Accurate identification of these subclones is therefore critical for reliable ploidy estimation. However, ploidy estimation based on a small number of cells (e.g., fewer than five cells) can be unreliable because of the substantial noise inherent in single-cell sequencing data. In contrast, ploidy estimates become more robust when derived from larger groups of cells sharing the same underlying ploidy state [49]. Accordingly, CNVeil performs hierarchical clustering with the objective of maximizing cluster size while maintaining ploidy homogeneity within each cluster.

Quantitatively, the minimum Ward linkage distance expected between tumor subclones with distinct ploidy states was empirically estimated from multiple real datasets (including T10, T16, and KTN302). This distance threshold, denoted by *D*_*p*_, is defined as *D*_*p*_ = 0.6 × *Height*_*Dendrogram*_, where *Height*_*Dendrogram*_ denotes the maximum height of the dendrogram generated during agglomerative clustering. Using *D*_*p*_ as the clustering cutoff, agglomerative clustering is applied to the tumor-cell subset of the matrix 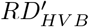, yielding candidate tumor subclones with consistent ploidy states. Subsequently, the numerical optimization procedure described above is applied independently to each subclone to estimate its optimal ploidy *ploidy*^*\**^, which is subsequently used for downstream total copy number inference.

#### Fine-grained clustering and across-cell breakpoint and segmentation identification

Cells within the same subclone generally share common copy number breakpoints and a common evolutionary history [50, 51]. Leveraging this shared structure can substantially improve breakpoint detection compared with analyzing each cell independently. To reduce the impact of cell-level noise and increase segmentation accuracy, CNVeil employs a cross-cell consensus breakpoint detection strategy that aggregates information across related cells. Before breakpoint detection, CNVeil further partitions each tumor subclone into finer cell clusters to maximize breakpoint homogeneity within clusters.

Similar to the previous stage of tumor subclone identification, an empirical threshold *D*_*b*_ is determined. This threshold represents the minimal Ward linkage distance between cell clusters groups with distinct breakpoint profiles. This threshold is estimated as *D*_*b*_ = 0.3 × *Height*_*Dendrogram*_, where *Height*_*Dendrogram*_ denotes the maximum height of the dendrogram generated during agglomerative clustering. Using *D*_*b*_ s the clustering cutoff, agglomerative clustering is applied to the tumor-cell subset of the matrix 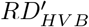 to generate finer clusters. Cells within each resulting cluster are expected to share a common set of copy number breakpoints.

Next, CNVeil performs cross-cell breakpoint detection independently within each cluster. To identify consensus breakpoints, CNVeil assesses the statistical significance of change points by aggregating read depth across cells within two target windows flanking each point and evaluating differences between windows across all change points. For each cell cluster, CNVeil takes the normalized read depth matrix as input, represented by a *p* × *m* matrix, where *p* denotes the number of cells in the cluster and *m* denotes the number of genomic bins. Given *m* bins, there are (*m* − 1) bin boundary points that can serve as candidate change points. We denote these (*m* − 1) points as the set *S*_*b*_. CNVeil then applies a sliding window of size *p* × *w*, where *w* is the window size (6 bins by default) to evaluate read depth difference across each candidate change points. Specifically, for the boundary point between *bin*_*k*_ and *bin*_*k*+1_, CNVeil calculates the median read depth within the left window, spanning *bin*_*max*(*k−w*+1,1)_ to *bin*_*k*_, denoted as *MRD*_*left*_ and the median read depth within the right window, spanning *bin*_*k*+1_ to *bin*_*min*(*k*+1+*w*,*m*)_, denoted as *MRD*_*right*_.

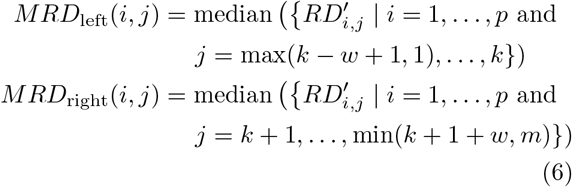

Next, CNVeil calculates the read-depth change rate for each candidate change point as:

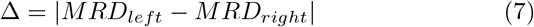

CNVeil computes Δ for each boundary point within each cell cluster, generating a set of read depth change rates denoted as *S*_Δ_. The changing rate threshold *t*_Δ_ is defined as the 90% quantile of *S*_Δ_. Any boundary point in *S*_*b*_ with a change rate exceeding the threshold *t*_Δ_ is designated and collected as a change point.

Within each cluster, a chaining algorithm is then applied to chain the change points into multiple disjoint regions. The chaining algorithm establishes an edge between any two change points if their distance is equal to one bin size, generating a network of multiple disjoint components. Each component forms a region that encompasses a real breakpoint in our assumption.

To control the false discovery rate of breakpoints, regions that contain less than six change points are excluded from the subsequent analysis since they are likely the artifact of sequencing noise. Finally, CNVeil selects the middle change point in each region as the predicted breakpoint. CNVeil repeats the cross-cell breakpoint detection procedure for each cell cluster and generates a unique breakpoint profile for each cluster.

Finally, genomics segmentation is constructed by defining each segment as the set of consecutive bins bounded by two adjacent predicted breakpoints. CNVeil also performs fine-grained clustering and across-cell breakpoints and segmentation identification for the initial normal cell cluster with the same approach.

#### Consensus read depth standardization and total copy number inference

In this module of the pipeline, the objective of CNVeil is to infer the copy number state for each bin of each cell. As we discussed before, for each cell, if we disregard the variances of read depth in each bin, the product of cell-specific ploidy and normalized read depth is the absolute copy number. To further normalize the read depth and minimize variance, CNVeil utilizes the cross-cell segmentation achieved in the previous module to standardize each normalized read depth 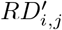 as below:

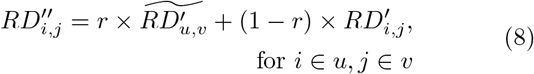

Where 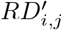 denotes the normalized read depth for the *i*-th cell and *j*-th bin, and 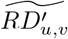 denotes the median read depth for the *v*-th segment in the *u*-th fine cell cluster. 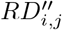 denotes the standardized read depth, and *r* is the standardization factor (0.5 by default). By standardizing, CNVeil ensures 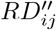 faithfully represents genuine genomic aberrations while minimizing the impact of artifact noise.

The Final Copy Number Profile (FCNP) is then inferred as:

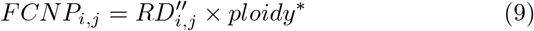

CNVeil uses subclone-level ploidy estimate, ploidy^*\**^, as a proxy for cell-specific ploidy because because cells within a subclone are assumed to share a common underlying ploidy state. Specifically, 1) ploidy^*\**^ is the optimal ploidy when maximizing the size of each cluster under the constraint that cells in one cluster share the same ploidy level; 2) Although normalized read-depth values may vary across cells because of technical and biological variability, the underlying ploidy is expected to remain largely consistent among cells belonging to the same subclone.

#### Allele-specific copy number inference

CNVeil infers allele-specific copy number by leveraging allelic imbalance signals from single-nucleotide polymorphisms (SNPs). Within each genomic segment, CNVeil assumes that the total copy number remains constant for a given cell. Under this assumption, an expectation-maximization (EM) algorithm is applied to the SNP-by-cell matrix to infer SNP phase assignments. Upon EM convergence, each SNP is assigned to one of two haplotypes, enabling estimation of cell-specific phased B-allele frequencies (BAFs) within each segment. Allele-specific copy number states are subsequently derived as the product of the phased BAF estimates and the total copy number of the corresponding segment.

#### Breakpoints pruning

Accurate segmentation is critical for reliable allele-specific copy number inference. To mitigate noise and reduce over-segmentation, CNVeil performs a multi-step smoothing and breakpoint pruning procedure. First, the total copy number profile (FCNP) matrix is smoothed using nonoverlapping 5Mb genomic bins, with values aggregated by the mean. CNVeil then refines the segmentation by identifying majority-supported breakpoints, denoted as *S*_*majorBND*_.

Over-segmentation often arises from copy number alterations present only in a small subset of cells, resulting in spurious breakpoints that do not reflect clonal events. To address this issue, CNVeil retains only breakpoints that are supported by at least 10% of tumor cells. After defining *S*_*majorBND*_, CNVeil recalculates the copy number for each cell by averaging values between consecutive major breakpoints. This procedure yields a simplified and denoised copy number matrix, substantially reducing model complexity and improving the robustness and computational efficiency of subsequent EM inference.

#### SNP extraction and filtering

Single-cell SNP information is extracted using Vatrix [52], generating a cell-by-SNP matrix in which each entry records the read support for reference and alternate alleles. SNPs are subsequently filtered according to their expected informativeness for allelic imbalance, with particularconsideration of loss-of-heterozygosity (LOH) scenarios.

In regions where both haplotypes are present, homozygous SNPs do not provide allelic imbalance information and are therefore excluded from EM algorithm. However, the interpretation of homozygous SNPs differs in LOH regions. In samples composed predominantly of tumor cells, LOH regions are expected to contain largely homozygous SNPs. In mixed tumor-normal samples, LOH regions are characterized by an elevated proportion of homozygous SNPs across cells. To account for these scenarios, CNVeil adopts a region-specific heuristic based on the proportion of homozygous SNPs. If the homozygous SNP fraction exceeds 0.9, the region is confidently classified as an LOH segment and EM inference is skipped. If the fraction lies between 0.6 and 0.9, indicating a mixture of LOH and non-LOH cells, SNPs are retained only when their aggregated read count exceeds 20. Finally, when the homozygous SNP fraction is below 0.6, suggesting little or no LOH, SNPs are filtered using both a BAF criterion (0.1-0.9) and aggregated read-count threshold (> 20) before EM inference.

#### EM algorithm

To infer allele-specific copy numbers, CNVeil maximizes a log-likelihood function

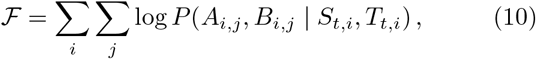

where *i* and *j* index cells and SNPs, respectively, within segment *t. A*_*i*,*j*_ and *B*_*i*,*j*_ denote the referenceand variant-supporting read counts for SNP *j* in cell *i. S*_*t*,*i*_ denotes the copy number of the major allele for cell *i* on segment *t*, and *T*_*t*,*i*_ is the corresponding total copy number from previous module.

Let *I*_*j*_ denote a latent indicator variable such that *I*_*j*_ = 1 if SNP *j* originates from the minor allele and *I*_*j*_ = 0 otherwise. The log-likelihood can be expressed as

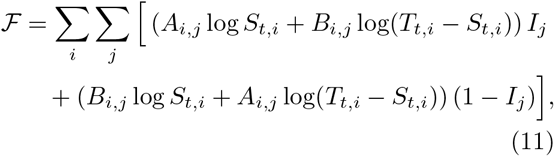

which explicitly constrains allele-specific copy numbers to sum to the inferred total copy number, preventing spurious allelic configurations driven by noisy BAF signals. More details on the derivation of Eqs. (10) and (11) are provided in the Supplementary Methods section.

CNVeil employs an Expectation-Maximization (EM) algorithm to optimize *F*. In the E-step, the posterior probability that SNP *j* originates from the minor allele is estimated as

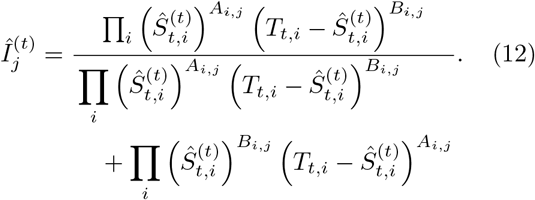

Where 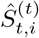 is the estimate of *S*_*t*,*i*_ at iteration *t*.

In the M-step, the major allele copy number is updated as

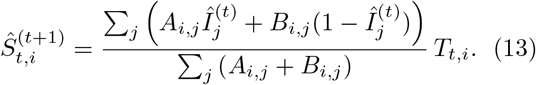

The EM procedure is iterated until convergence, after which *Ŝ*_*t*,*i*_ is rounded to the nearest integer to ensure integer-valued copy number estimate. The algorithm is initialized with 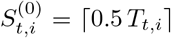, corresponding to an equal allocation of the total copy number between the two alleles in the absence of prior evidence for allelic imbalance.

#### Haplotype-specific copy number reconstruction

The expectation-maximization algorithm described above infers allele-specific copy number independently for each segment. Consequently, the inferred allelespecific copy number states represent an unordered pair of alleles and do not preserve chromosome-wide parental haplotype identity. To reconstruct haplotype-specific copy number profiles, CNVeil assigns a consistent parental orientation across adjacent segments using a dynamic programming algorithm following the framework introduced in CHISEL [27].

Following the dynamic programming framework introduced in CHISEL [27], chromosome-wide haplotype reconstruction is formulated as an optimization problem over segment orientations. Let

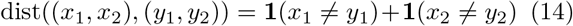

denote the Hamming distance between two haplotype-specific copy number states. The transition cost between adjacent genomic segments is then defined as

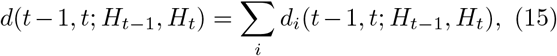

The optimal chromosome-wide orientation sequence is obtained by minimizing the cumulative transition cost,

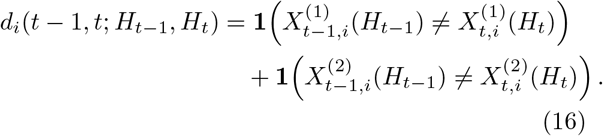

Because the objective depends only on adjacent genomic segments, the global optimum can be obtained exactly using dynamic programming. Let *D*(*t, H*_*t*_) denote the minimum cumulative transition cost for the first *t* genomic segments ending in orientation *H*_*t*_. The recurrence relation is

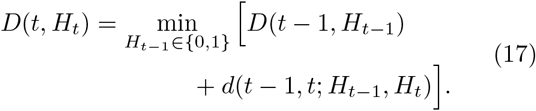

with initialization *D*(1, 0) = *D*(1, 1) = 0. The optimal orientation sequence is recovered by backtracking from the terminal state with the minimum cumulative transition cost. Applying the inferred orientations to the unordered allele-specific copy number states yields chromosome-scale haplotype-specific copy number profiles.

### Simulated data by simSCSnTree

We employed SimSCSnTree to generate simulated datasets with a gold standard. The simulation process executed by SimSCSnTree is intricately designed in three primary steps, reflecting the complexities inherent in cancer genomics.

#### Step 1: Phylogenetic tree construction and genomic variation modeling

In the initial step, a phylogenetic tree is crafted using SimSCSnTree, with each node representing a distinct genome segment, and the connecting edges denoting genetic variations (CNVs). Built upon the hg19 reference genome, it ensures our simulation mirrors actual genomic structures and sequences. The outcome produces two critical numpy (.npy) files: one delineating the CNVs alongside the tree structure and another storing intermediate information essential for the subsequent simulation step.

SimSCSnTree controls the characteristics of the tree and the final CNV profile by multiple parameters, such as the multiplier of the mean CNV on the root, the rate of deletion, the mean, and median of the tree width distribution, etc. For example, to generate the tree for a dataset with an average ploidy of 1.5, we used the command below:

~~~
python ./SimSCSnTree/main.par.overlapping.py \
-S ./SimSCSnTree/wgsim-master \
-r ./simulate_ploidy1.5 \
-t reference_hg19.fa \
-n 100 -p 1 -X 4 -W 0 -m 5000000 -e 20000000 \
-d 1 -c 8 -E 1 -F 8 -H 0.000001
~~~

where **-n** is number of cells, **-p** is number of processors, **-X** is the multiplier of the mean CNV on root, **-W** denotes whether there is whole chromosome amplification (0 indicating no), **-m** is the minimum copy number size, **-e** is the parameter for the exponential distribution for copy number size beyond the minimum one, **-d** is the rate of deletion, **-c** is the average number of copy number variations to be added on a branch, **-E** is the whole amplification copy number addition, **-F** is the mean of the tree width distribution, and **-H** is the standard deviation of the tree width distribution.

To generate a dataset of other ploidy, the user needs to adjust the corresponding parameters. To reproduce all four simulated datasets in this paper, users can refer to our GitHub for more details.

#### Step 2: Read sampling from the simulated genome building upon the tree

In the second step, reads are sampled from the genome at selected nodes of the tree. This phase is critical in translating theoretical genomic information into practical sequencing data. By utilizing the .npy files generated from the first step, this phase simulated the sequencing process, closely resembling real-world sequencing techniques.

For example, to simulate reads from the tree generated by step1, we used the command below:

~~~
python ./SimSCSnTree/main.par.overlapping.py \
-p 10 -k 1 \
-r ./simulate_ploidy1.5/ \
-S ./SimSCSnTree/wgsim-master/ \
--template-ref genome_hg19.fa \
--Lorenz-y 0.28 -n 100 -L -1 -Y 0.8 -v 0.01 -l 70
~~~

where **-p** is the number of processors, **-k** denotes whether to skip step 1 (1 for skip), **–Lorenz-y** is the value on the y-axis of the Lorenz curve, **-n** is number of cells, **-l** is the read length, **-v** is the average coverage of the sequence, **-L** controls the levels to be sequenced, and **-Y** specifies a range of nodes to sequence. the above command for step 2 was employed for generating all four simulated datasets in this paper.

#### Step 3: Extract the gold standard set

The ground truth data for CNVs obtained from this simulation served as the gold standard against which the accuracy and effectiveness of various CNV inference tools can be assessed. The ground truth was extracted from different nodes (tumor subclones), that contain a set of breakpoints (where the copy number changes) and the left and right copy numbers for each breakpoint. Similarly, the callset from each CNV inference tool was also reformatted into a set of breakpoints and neighboring copy numbers for a fair comparison and evaluation.

#### Evaluation against gold standard in simulated datasets

In simulated data, we compared the callset by each tool with the gold standard set using two evaluation modes: segmentation mode and CNV mode.

In segmentation evaluation mode, we required the position of the called breakpoint and gold standard breakpoint to approximately align. The called breakpoint matches the gold standard breakpoint if they satisfy the below conditions:

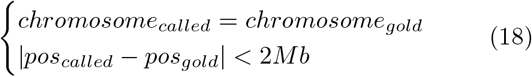

In stringent CNV evaluation mode, we required both the breakpoints and the copy number on the left and right bins of the breakpoint to match between the called event and the gold standard. The called event matches the gold standard if they satisfy the below conditions:

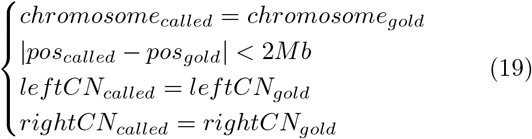

Following an N-to-N matching procedure between each callset and the gold standard set, breakpoints from the callset matched to any breakpoint in the gold standard set are classified as true positives (TPs), while the remaining breakpoints from the callset are considered false positives (FPs). Conversely, breakpoints from the gold standard set without a matching counterpart in the callset are classified as false negatives (FNs). The evaluation matrices are calculated as below:

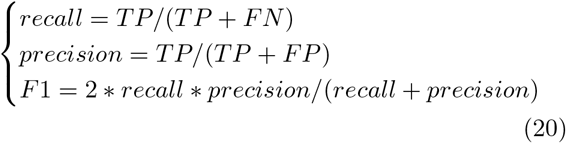

### Validation of allele-specific copy number states using aggregated BAF

We generated aggregated B-allele frequency (BAF) histograms to evaluate whether inferred allele-specific copy number states were supported by SNP allele counts.

First, copy number segments were constructed from the allele-specific copy number intervals inferred by CNVeil. For each cell *i* and bin *j*, the inferred allelespecific copy number state was denoted as *A*_*ij*_ *B*_*ij*_, and total copy number was calculated as

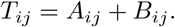

Adjacent bins were merged unless a total-copy-number change was observed across the boundary. Specifically, for each boundary between bins *j* and *j*+1, we counted the number of tumor cells with *T*_*ij*_ *≠ T*_*i*,*j*+1_. Boundaries supported by more than 10% of tumor cells were retained and used to define the final segments. These segments inferred from CNVeil result were later applied to other methods for consistency.

Second, cells within each segment were grouped based on the allele-specific copy number profiles inferred by CNVeil. Cells were assigned to the same group if they had identical *A*|*B* profiles across all bins within the segment. Cell groups containing fewer than 100 cells were excluded from downstream analysis. This cell grouping was later applied to other method for BAF calculation.

Thirdly, for each tool, SNP-level allele counts were aggregated within each retained cell group and analysis segment inferred from CNVeil result. For each SNP position *k*, reference and alternate allele counts were summed across all cells in the group:

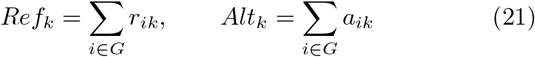

where *G* denotes the cell group, and *r*_*ik*_ and *a*_*ik*_ are the reference and alternate allele counts for cell *i* at SNP *k*, respectively. The aggregated BAF for SNP *k* was then calculated as:

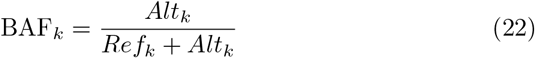

Only SNPs with aggregated depth *Ref*_*k*_ + *Alt*_*k*_ > 10 were retained.

Finally, BAF values were visualized as histograms with 50 bins between 0 and 1. For comparison, the expected BAF values implied by an inferred allelespecific copy number state *A*|*B* were calculated as:

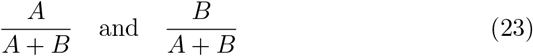

and overlaid as dashed vertical lines. Agreement between the observed BAF histogram peaks and the expected BAF values was used to visually assess support for the inferred allele-specific copy number state.

### Comparison of pseudo-bulk and matched bulk WES copy number profiles

To evaluate the accuracy of CNVeil, we compared pseudo-bulk haplotype-specific copy number profiles derived from single-cell DNA sequencing with matched bulk copy number profiles generated by ASCAT. For each sample, haplotype-specific copy number states were averaged across cells to generate pseudo-bulk profiles using 5-Mb genomic bins. ASCAT segments were mapped to the same genomic regions based on overlap between ASCAT segments and pseudo-bulk bins, and the corresponding major and minor copy number states were assigned to each bin.

Because ASCAT reports unordered major and minor copy numbers whereas CNVeil infers haplotype-specific copy number states, the two profiles were aligned before comparison. For each segment, we calculated the direction of allelic imbalance in the bulk and pseudo-bulk profiles as

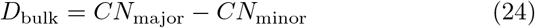

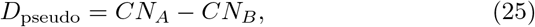

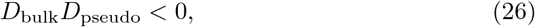

The direction of allelic imbalance was considered inconsistent between the two profiles, and the ASCAT major and minor copy number states were exchanged:

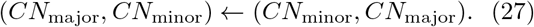

After alignment, concordance between ASCAT and CNVeil was evaluated using Pearson correlation coefficients for haplotype-specific and total copy number profiles. Agreement of discrete copy number states was assessed using copy number state confusion matrices.

## Code availability

All code is available at https://github.com/maiziezhoulab/CNVeil.

## Acknowledgements

This work was supported by the NIH NIGMS Maximizing Investigators’ Research Award (MIRA) R35 GM146960 (X.M.Z), Waddell Walker Hancock Cancer Discovery Fund Award (X.M.Z), and NSF CCF grant number 2523717 (X.M.).

## Author’s contributions

X.M.Z. and X.M. conceived and led this work. W.Y., C.L, X.M., and X.M.Z. designed the framework and W.Y. and C.L implemented the framework. W.Y., C.L., Y.H., L.Z., Z.W., Y.H.L performed data analysis. L.Z. assisted with the simulation analysis. W.Y., C.L, X.M., and X.M.Z. wrote the manuscript with input from all authors.

