## Supplementary for "CNVeil resolves haplotype-specific copy number and uncovers subclonal architecture hidden from total copy number profiling in single-cell cancer genomes"

Yuan et al.

**References** **19**

### S1 Supplementary Results

#### S1.1 Evaluation on the scDNA-seq data of triple-negative cancer patient KTN302

We further applied the analysis to the scDNA-seq data of a triple-negative breast cancer patient, KTN302, whose treatment stages were well-documented [1]. This dataset includes 47 cells from the pre-treatment stage and 45 cells from the mid-treatment stage (**Table 2**).

To again ensure a fair and transparent comparison, we systematically ordered and organized cells in heatmaps based on the pre-treatment and mid-treatment stages (**Supplementary Fig. 2a**). We observed that CNVeil successfully detected a normal cell population from the mid-treatment stage and two subclones of aneuploid cells in the pre-treatment tumors. Although SCOPE showed a chaotic CNV profile at the mid-treatment stage, it was able to differentiate the tumor subclone from the normal cell population. Nevertheless, it is crucial to note that the KTN302 data is highly noisy. Therefore, SCOPE had to utilize mid-stage information as prior knowledge to select normal cells, achieving a reasonable outcome. When stage information was not provided to SCOPE, its CNV profile in the mid-treatment stage revealed many incorrect hyperdiploid cells (Figure S8). SeCNV, AneuFinder, and Ginkgo all exhibited numerous inaccuracies, including both hyperdiploid and hypodiploid cells during the mid-treatment stage, along with a few hypodiploid cells in the pre-treatment stage. These results indicated that when the scDNA-seq data was highly noisy, existing tools often failed to identify normal cells and correct bias, or they needed prior knowledge to assist these steps. SEACON systematically inferred elevated copy number states across most cells, whereas FLCNA shifted much of the genome toward lower copy number states. Alleloscope failed to recover the overall copy number landscape regardless of whether the top candidate diploid regions or the consensus diploid regions were used for normalization, assigning total copy number 2 to the majority of genomic segments and thereby masking much of the underlying subclonal copy number variation. HMMcopy tended to overestimate the ploidy for the majority of cells in the mid-treatment stages. CNRein broadly preserved the chromosome-scale copy number landscape but substantially overestimated ploidy in the mid-treatment normal-cell population, with many normal cells assigned copy number states of 5 or 6 across large genomic regions. whereas CHISEL generated substantially noisier allele-derived copy number profiles throughout the genome.

#### S1.2 Evaluation on the scDNA-seq data of breast cancer patients T16

We also extended the analysis to another breast cancer patient, identified as T16 [2]. This dataset comprises 52 cells from primary tumor sites and 48 cells from metastatic tumor sites (**Table 2**). Both sites include a mixture of tumor and normal cells. Compared to T10, T16 demonstrated more homogeneous tumor subclones in both primary tumor and metastasis, indicating a monogenomic tumor.

The heatmaps in (**Supplementary Fig. 2b**) reveal that most tools demonstrated similar characteristics as what we observed in T10. Overall, CNVeil, SeCNV, SCOPE, SPRINTER, AneuFinder, and Ginkgo showed a generally similar CNV profile pattern, distinguishing between a normal cell subclone and one subclone of hyperdiploid cancer cells. However, SeCNV, SCOPE, AneuFinder, and Ginkgo exhibited a few cells with incorrect ploidy and outlier bins across the genome. SEACON systematically overestimated copy number states across much of the genome, whereas FLCNA underestimated copy number states throughout the majority of cells. Alleloscope failed to recover the overall copy number landscape, with most genomic segments assigned intermediate copy number states and reduced subclonal resolution. Upon careful inspection, CNVeil, SeCNV, SCOPE, AneuFinder, and Ginkgo all successfully identified two tumor subclones: primary and metastasis subclones (trisomic state). However, there were some disagreements regarding the copy number, especially on chromosomes 4, 5, and 7. CNVeil, SeCNV, AneuFinder, and Ginkgo predominantly predicted a copy number of 3 or 4 for most bins on chromosomes 4 and 5, whereas SCOPE predicted a copy number of 2. Similarly, for chromosome 7, CNVeil, SeCNV, AneuFinder, and Ginkgo predicted a copy number of 7, whereas SCOPE predicted a copy number of 5 or 6. In general, SCOPE tended to underestimate the copy numbers compared to CNVeil, SeCNV, and AneuFinder on T16 tumor cells. Conversely, the heatmap by HMMcopy showed a separation between normal and tumor cells, but frequently assigned copy number 1 to diploid genomic regions, resulting in more homogeneous copy number profiles across cells. CNRein broadly preserved the chromosome-scale copy number landscape but exhibited noisier segmentation across the genome. CHISEL showed a chaotic CNV profile, where there was no clear separation between normal and tumor cells. In summary, CNVeil still demonstrated the best CNV profiles for subclone identification and segmentation.

### S2 Supplementary Methods

#### S2.1 Allele-specific copy number inference

**Notation and setup.** Let $i \in \{1, \ldots, n\}$ index cells and $j \in \{1, \ldots, J_r\}$ index phased heterozygous single-nucleotide variants (SNVs) located within genomic segment $r$. For each cell-SNV pair, let $A_{ij}$ and $B_{ij}$ denote the reference- and alternative-supporting read counts, respectively, with total coverage $N_{ij} = A_{ij} + B_{ij}$.

Let $T_{ir} \in \mathbb{Z}_{\geq 0}$ denote the total copy number of segment $r$ in cell $i$, inferred from the final copy number profile (FCNP). We denote by $S_{ir} \in \{0, 1, \ldots, T_{ir}\}$ the copy number of the major allele in segment $r$ for cell $i$, and the corresponding minor-allele copy number is $T_{ir} - S_{ir}$.

To model the unknown allelic origin of each SNV, we introduce a latent indicator variable

$$I_j = \begin{cases} 1, & \text{if SNV } j \text{ resides on the minor allele,} \\ 0, & \text{if SNV } j \text{ resides on the major allele.} \end{cases}$$

In the absence of prior allelic information, a symmetric Bernoulli prior is assumed,

$$P(I_j = 1) = P(I_j = 0) = \tfrac{1}{2}.$$

**Observation model.** Conditioned on the allelic origin of an SNV and on the underlying copy number configuration, the observed read counts are modeled using a binomial sampling process. If SNV $j$ resides on the minor allele ($I_j = 1$),

$$A_{ij} \mid I_j = 1, S_{ir}, T_{ir} \sim \text{Binomial}\left(N_{ij}, \frac{S_{ir}}{T_{ir}}\right),$$

whereas if SNV $j$ resides on the major allele ($I_j = 0$),

$$A_{ij} \mid I_j = 0, S_{ir}, T_{ir} \sim \text{Binomial}\left(N_{ij}, \frac{T_{ir} - S_{ir}}{T_{ir}}\right).$$

Up to multiplicative constants independent of $S_{ir}$, the joint likelihood of observing $(A_{ij}, B_{ij})$ is therefore

$$P(A_{ij}, B_{ij} \mid I_j, S_{ir}, T_{ir}) \propto \begin{cases} S_{ir}^{A_{ij}} (T_{ir} - S_{ir})^{B_{ij}}, & I_j = 0, \\ (T_{ir} - S_{ir})^{A_{ij}} S_{ir}^{B_{ij}}, & I_j = 1. \end{cases}$$

**Complete-data log-likelihood.** Let $\mathbf{I} = \{I_j\}$ and $\mathbf{S}_r = \{S_{ir}\}$. The complete-data log-likelihood for segment $r$ is

$$\mathcal{F}(\mathbf{S}_r; \mathbf{I}) = \sum_{i=1}^{n} \sum_{j=1}^{J_r} \Big[ I_j \big( A_{ij} \log S_{ir} + B_{ij} \log(T_{ir} - S_{ir}) \big) + (1 - I_j) \big( A_{ij} \log(T_{ir} - S_{ir}) + B_{ij} \log S_{ir} \big) \Big] + C,$$

where $C$ collects terms independent of $S_{ir}$.

**Expectation–Maximization algorithm.** **E-step.** Given current estimates $\hat{S}_{ir}^{(t)}$, the posterior expectation $\gamma_j^{(t)} = \mathbb{E}[I_j \mid \cdot]$ is computed as

$$\gamma_j^{(t)} = \frac{\prod_i (\hat{S}_{ir}^{(t)})^{A_{ij}} (T_{ir} - \hat{S}_{ir}^{(t)})^{B_{ij}}}{\prod_i (\hat{S}_{ir}^{(t)})^{A_{ij}} (T_{ir} - \hat{S}_{ir}^{(t)})^{B_{ij}} + \prod_i (\hat{S}_{ir}^{(t)})^{B_{ij}} (T_{ir} - \hat{S}_{ir}^{(t)})^{A_{ij}}}.$$

For numerical stability, the posterior is evaluated in log-space. Define

$$\Delta_j^{(t)} = \sum_i (A_{ij} - B_{ij}) \left[ \log \hat{S}_{ir}^{(t)} - \log(T_{ir} - \hat{S}_{ir}^{(t)}) \right],$$

so that

$$\gamma_j^{(t)} = \frac{1}{1 + \exp(-\Delta_j^{(t)})}.$$

**M-step.** Define the weighted sufficient statistics

$$U_{ir}^{(t)} = \sum_j \left( A_{ij}\gamma_j^{(t)} + B_{ij}(1 - \gamma_j^{(t)}) \right),$$
$$V_{ir}^{(t)} = \sum_j \left( B_{ij}\gamma_j^{(t)} + A_{ij}(1 - \gamma_j^{(t)}) \right).$$

Since

$$U_{ir}^{(t)} + V_{ir}^{(t)} = \sum_j (A_{ij} + B_{ij}),$$

The update for the major-allele copy number is

$$\hat{S}_{ir}^{(t+1)} = \frac{U_{ir}^{(t)}}{U_{ir}^{(t)} + V_{ir}^{(t)}} T_{ir}.$$

To enforce integer copy numbers, the estimate is projected onto $\{0, 1, \ldots, T_{ir}\}$ by rounding. Initialization is performed using

$$S_{ir}^{(0)} = \left\lceil \tfrac{1}{2} T_{ir} \right\rceil.$$

#### S2.2 Evolutionary analysis from haplotype-resolved copy number profiles

To investigate tumor evolution at haplotype resolution, haplotype-specific copy number profiles inferred by CNVeil were used for subclone identification, consensus profile generation, phylogenetic reconstruction, and downstream characterization of lineage-specific copy number alterations.

##### S2.2.1 Haplotype-based subclone inference

For each cell $i$, the genome was represented as a sequence of haplotype-specific copy number states across $M$ genomic bins,

$$\mathbf{h}_i = \{(a_{i1}, b_{i1}), (a_{i2}, b_{i2}), \ldots, (a_{iM}, b_{iM})\}, \tag{1}$$

where $a_{im}$ and $b_{im}$ denote the inferred copy numbers of the two haplotypes in genomic bin $m$.

Pairwise distances between cells were computed as the fraction of genomic bins with discordant haplotype-specific copy number states,

$$d(i, j) = \frac{1}{M} \sum_{m=1}^{M} \mathbb{I}\left[(a_{im}, b_{im}) \neq (a_{jm}, b_{jm})\right], \tag{2}$$

where $\mathbb{I}(\cdot)$ denotes the indicator function. Weighted hierarchical clustering was then performed on the resulting distance matrix to identify haplotype-defined subclones.

###### S2.2.2 Consensus copy number profile generation

For each inferred subclone $C_k$, consensus haplotype-specific copy number was calculated as the median across all cells,

$$\hat{a}_{km} = \text{median}_{i \in C_k}(a_{im}), \qquad \hat{b}_{km} = \text{median}_{i \in C_k}(b_{im}), \tag{3}$$

where $\hat{a}_{km}$ and $\hat{b}_{km}$ denote the consensus copy numbers of the two haplotypes for genomic bin $m$ in subclone $k$. The corresponding consensus total copy number was calculated as

$$\hat{t}_{km} = \hat{a}_{km} + \hat{b}_{km}. \tag{4}$$

Consensus profiles were used for all downstream analyses.

###### S2.2.3 Phylogenetic reconstruction

Consensus haplotype-specific copy number profiles were analyzed using MEDICC2 [3]. Haplotype-resolved phylogenies were reconstructed using chromosome-wise bootstrapping, whereas total copy number trees were reconstructed from the corresponding consensus total copy number profiles.

The haplotype-resolved trees were generated using

```
medicc2 input_medicc2.tsv ./ \
    --plot auto \
    --events \
    --bootstrap-method chr-wise \
    --bootstrap-nr 100 \
    -j 10
```

For comparison, total copy number trees were generated using

```
medicc2 subclone_totalcn.tsv ./ \
    --total-copy-numbers \
    --input-allele-columns total_cn \
    --plot auto \
    --events \
    --bootstrap-method chr-wise \
    --bootstrap-nr 100 \
    -j 10
```

The branch-specific copy number events inferred by MEDICC2 were subsequently used for downstream gene annotation.

###### S2.2.4 Gene annotation and copy number event classification

Consensus genomic bins were intersected with RefSeq gene annotations. A genomic bin $B_m = [s_m, e_m]$ was considered to overlap a gene interval $G = [s_G, e_G]$ when

$$s_m < e_G \qquad \text{and} \qquad e_m > s_G. \tag{5}$$

Each genomic bin was assigned one event label,

$$E_{km} \in \{\text{Amp}, \text{Del}, \text{LOH}, \text{CN-LOH}, \text{AI}, \text{Neutral}\},$$

according to the consensus haplotype-specific copy number state.

Amplification was defined as

$$\hat{a}_{km} + \hat{b}_{km} > 2. \tag{6}$$

Deletion was defined as

$$\hat{a}_{km} + \hat{b}_{km} < 2. \tag{7}$$

Loss of heterozygosity (LOH) was defined as

$$\min(\hat{a}_{km}, \hat{b}_{km}) = 0, \qquad \hat{a}_{km} + \hat{b}_{km} > 2. \tag{8}$$

Copy-neutral loss of heterozygosity (CN-LOH) was defined as

$$\min(\hat{a}_{km}, \hat{b}_{km}) = 0, \qquad \hat{a}_{km} + \hat{b}_{km} = 2. \tag{9}$$

Allelic imbalance (AI) was defined as

$$\hat{a}_{km} \neq \hat{b}_{km}, \qquad \min(\hat{a}_{km}, \hat{b}_{km}) > 0. \tag{10}$$

##### S2.2.5 Gene-level event frequency

For each gene $g$, the frequency of copy number alteration across inferred subclones was calculated as

$$f_g = \frac{1}{K} \sum_{k=1}^{K} \mathbb{I}\left[\exists\, m \in \mathcal{B}(g) : (\hat{a}_{km}, \hat{b}_{km}) \neq (1, 1)\right], \tag{11}$$

where $K$ denotes the number of inferred subclones and $\mathcal{B}(g)$ denotes the set of genomic bins overlapping gene $g$.

Similarly, the frequency of each event type $e$ was calculated as

$$f_{g,e} = \frac{1}{K} \sum_{k=1}^{K} \mathbb{I}\left[\exists\, m \in \mathcal{B}(g) : E_{km} = e\right], \tag{12}$$

where $E_{km}$ denotes the event assigned to genomic bin $m$ in subclone $k$.

### S3 Supplementary Tables

| Ploidy | Tool | F1 | Precision | Recall | $n$ |
|---|---|---|---|---|---|
| 1.5 | CNVeil | **0.7817 ± 0.0272** [0.7830] | **0.8565 ± 0.0293** [0.8600] | **0.7191 ± 0.0284** [0.7155] | 95 |
| | Ginkgo | 0.6042 ± 0.3300 [0.7789] | 0.6648 ± 0.3636 [0.8542] | 0.5540 ± 0.3026 [0.7115] | 95 |
| | AneuFinder | 0.6754 ± 0.0272 [0.6780] | 0.6599 ± 0.0351 [0.6667] | 0.6925 ± 0.0270 [0.6949] | 95 |
| | SeCNV | 0.5791 ± 0.3109 [0.7200] | 0.6661 ± 0.3536 [0.8454] | 0.5140 ± 0.2801 [0.6364] | 95 |
| | SCOPE | 0.0053 ± 0.0059 [0.0059] | 0.0041 ± 0.0046 [0.0045] | 0.0076 ± 0.0083 [0.0086] | 51 |
| | SPRINTER | 0.4400 ± 0.1679 [0.4545] | 0.5969 ± 0.2210 [0.6393] | 0.3492 ± 0.1366 [0.3558] | 95 |
| | HMMcopy | 0.0000 ± 0.0000 [0.0000] | 0.0000 ± 0.0000 [0.0000] | 0.0000 ± 0.0000 [0.0000] | 51 |
| 3 | CNVeil | **0.6964 ± 0.0382** [0.7030] | **0.7284 ± 0.0417** [0.7377] | **0.6682 ± 0.0449** [0.6628] | 97 |
| | Ginkgo | 0.4513 ± 0.0886 [0.4474] | 0.5169 ± 0.0994 [0.5294] | 0.4024 ± 0.0855 [0.3953] | 97 |
| | AneuFinder | 0.3283 ± 0.0445 [0.3313] | 0.2955 ± 0.0407 [0.3000] | 0.3714 ± 0.0566 [0.3636] | 97 |
| | SeCNV | 0.3029 ± 0.0428 [0.3000] | 0.5159 ± 0.0645 [0.5000] | 0.2160 ± 0.0363 [0.2209] | 97 |
| | SCOPE | 0.5725 ± 0.0699 [0.5714] | 0.6296 ± 0.0815 [0.6294] | 0.5265 ± 0.0680 [0.5250] | 42 |
| | SPRINTER | 0.2119 ± 0.0851 [0.1920] | 0.3507 ± 0.1094 [0.3256] | 0.1539 ± 0.0694 [0.1395] | 97 |
| | HMMcopy | 0.0527 ± 0.0280 [0.0515] | 0.1181 ± 0.0665 [0.1133] | 0.0342 ± 0.0180 [0.0341] | 42 |
| 4 | CNVeil | **0.4804 ± 0.0755** [0.4881] | **0.5087 ± 0.0956** [0.5111] | **0.4578 ± 0.0622** [0.4741] | 95 |
| | Ginkgo | 0.3033 ± 0.0744 [0.3141] | 0.3674 ± 0.0957 [0.3902] | 0.2592 ± 0.0616 [0.2683] | 95 |
| | AneuFinder | 0.2582 ± 0.0558 [0.2559] | 0.2554 ± 0.0634 [0.2545] | 0.2632 ± 0.0505 [0.2561] | 95 |
| | SeCNV | 0.2355 ± 0.0529 [0.2436] | 0.4556 ± 0.0997 [0.4750] | 0.1589 ± 0.0363 [0.1638] | 95 |
| | SCOPE | 0.2459 ± 0.0541 [0.2514] | 0.2589 ± 0.0697 [0.2673] | 0.2362 ± 0.0422 [0.2439] | 44 |
| | SPRINTER | 0.0028 ± 0.0058 [0.0000] | 0.0039 ± 0.0082 [0.0000] | 0.0021 ± 0.0045 [0.0000] | 95 |
| | HMMcopy | 0.1572 ± 0.0439 [0.1646] | 0.3423 ± 0.1101 [0.3442] | 0.1025 ± 0.0283 [0.1034] | 44 |
| 5 | CNVeil | **0.6750 ± 0.1099** [0.7175] | **0.7381 ± 0.1051** [0.7725] | **0.6245 ± 0.1119** [0.6668] | 96 |
| | Ginkgo | 0.2048 ± 0.1503 [0.2810] | 0.2680 ± 0.1969 [0.3599] | 0.1661 ± 0.1223 [0.2244] | 96 |
| | AneuFinder | 0.2167 ± 0.0862 [0.2376] | 0.2266 ± 0.0906 [0.2512] | 0.2088 ± 0.0848 [0.2266] | 96 |
| | SeCNV | 0.0029 ± 0.0142 [0.0000] | 0.0071 ± 0.0322 [0.0000] | 0.0019 ± 0.0091 [0.0000] | 96 |
| | SCOPE | 0.0123 ± 0.0117 [0.0087] | 0.0123 ± 0.0117 [0.0087] | 0.0123 ± 0.0118 [0.0093] | 45 |
| | SPRINTER | 0.0005 ± 0.0023 [0.0000] | 0.0010 ± 0.0047 [0.0000] | 0.0003 ± 0.0015 [0.0000] | 96 |
| | HMMcopy | 0.0000 ± 0.0000 [0.0000] | 0.0000 ± 0.0000 [0.0000] | 0.0000 ± 0.0000 [0.0000] | 45 |

Table S1: CNV-level performance of total copy number state identification across simulated ploidy settings. F1 score, precision, and recall are reported for simulated datasets with average ploidy levels of 1.5, 3, 4, and 5. Values are presented as mean ± standard deviation (s.d.), with the median shown in brackets. $n$ denotes the number of successfully evaluated datasets for each method after excluding failed runs. The best-performing method (highest mean value) for each metric within each ploidy level is shown in **bold**.

| Ploidy | Tool | F1 | Precision | Recall | $n$ |
|---|---|---|---|---|---|
| 1.5 | CNVeil | **0.8367 ± 0.0247** [0.8454] | 0.9169 ± 0.0285 [0.9184] | 0.7697 ± 0.0255 [0.7759] | 95 |
| | Ginkgo | 0.8239 ± 0.0211 [0.8263] | 0.9048 ± 0.0299 [0.9080] | 0.7566 ± 0.0219 [0.7586] | 95 |
| | SCOPE | 0.5475 ± 0.0406 [0.5595] | 0.4208 ± 0.0390 [0.4299] | **0.7854 ± 0.0348** [0.8017] | 51 |
| | AneuFinder | 0.7284 ± 0.0274 [0.7305] | 0.7118 ± 0.0384 [0.7167] | 0.7467 ± 0.0238 [0.7414] | 95 |
| | SeCNV | 0.7893 ± 0.0655 [0.8182] | 0.8941 ± 0.0396 [0.8953] | 0.7094 ± 0.0838 [0.7672] | 95 |
| | HMMcopy | 0.7219 ± 0.0349 [0.7191] | **0.9205 ± 0.0223** [0.9200] | 0.5946 ± 0.0409 [0.5865] | 51 |
| | SPRINTER | 0.5039 ± 0.0941 [0.4804] | 0.6843 ± 0.1156 [0.6774] | 0.3997 ± 0.0808 [0.3818] | 95 |
| 3 | CNVeil | **0.7993 ± 0.0343** [0.8032] | **0.8365 ± 0.0478** [0.8442] | **0.7665 ± 0.0364** [0.7727] | 97 |
| | Ginkgo | 0.5939 ± 0.0508 [0.5974] | 0.6808 ± 0.0599 [0.6842] | 0.5292 ± 0.0573 [0.5250] | 97 |
| | SCOPE | 0.6716 ± 0.0527 [0.6795] | 0.7385 ± 0.0659 [0.7500] | 0.6176 ± 0.0548 [0.6163] | 42 |
| | AneuFinder | 0.5027 ± 0.0444 [0.5034] | 0.4532 ± 0.0477 [0.4490] | 0.5676 ± 0.0542 [0.5698] | 97 |
| | SeCNV | 0.4293 ± 0.0464 [0.4427] | 0.7303 ± 0.0502 [0.7368] | 0.3063 ± 0.0454 [0.3182] | 97 |
| | HMMcopy | 0.3147 ± 0.0632 [0.3286] | 0.7010 ± 0.0848 [0.7042] | 0.2055 ± 0.0494 [0.2107] | 42 |
| | SPRINTER | 0.3090 ± 0.1037 [0.2992] | 0.5124 ± 0.1322 [0.5143] | 0.2241 ± 0.0852 [0.2125] | 97 |
| 4 | CNVeil | **0.6418 ± 0.0687** [0.6495] | 0.6792 ± 0.0981 [0.7033] | **0.6120 ± 0.0503** [0.6293] | 95 |
| | Ginkgo | 0.5249 ± 0.0603 [0.5376] | 0.6346 ± 0.0895 [0.6667] | 0.4492 ± 0.0473 [0.4528] | 95 |
| | SCOPE | 0.3994 ± 0.0546 [0.4087] | 0.4193 ± 0.0822 [0.4279] | 0.3847 ± 0.0312 [0.3932] | 44 |
| | AneuFinder | 0.4489 ± 0.0489 [0.4464] | 0.4431 ± 0.0668 [0.4563] | 0.4585 ± 0.0369 [0.4569] | 95 |
| | SeCNV | 0.3926 ± 0.0437 [0.4103] | **0.7610 ± 0.0908** [0.7949] | 0.2648 ± 0.0298 [0.2759] | 95 |
| | HMMcopy | 0.3434 ± 0.0425 [0.3447] | 0.7386 ± 0.0894 [0.7588] | 0.2245 ± 0.0310 [0.2241] | 44 |
| | SPRINTER | 0.3096 ± 0.0523 [0.3174] | 0.4719 ± 0.0960 [0.4915] | 0.2330 ± 0.0426 [0.2333] | 95 |
| 5 | CNVeil | **0.8152 ± 0.0837** [0.8474] | **0.8929 ± 0.0711** [0.9131] | **0.7537 ± 0.0954** [0.7812] | 96 |
| | Ginkgo | 0.4960 ± 0.0480 [0.4988] | 0.6699 ± 0.0587 [0.6667] | 0.3961 ± 0.0477 [0.3984] | 96 |
| | SCOPE | 0.5373 ± 0.0454 [0.5447] | 0.5539 ± 0.0639 [0.5538] | 0.5261 ± 0.0448 [0.5391] | 45 |
| | AneuFinder | 0.4478 ± 0.0439 [0.4526] | 0.4745 ± 0.0413 [0.4757] | 0.4272 ± 0.0561 [0.4310] | 96 |
| | SeCNV | 0.2558 ± 0.0377 [0.2482] | 0.7708 ± 0.0757 [0.7727] | 0.1549 ± 0.0290 [0.1441] | 96 |
| | HMMcopy | 0.3285 ± 0.0673 [0.3270] | 0.7870 ± 0.0768 [0.7931] | 0.2094 ± 0.0484 [0.2037] | 45 |
| | SPRINTER | 0.2696 ± 0.0531 [0.2695] | 0.5348 ± 0.0890 [0.5432] | 0.1824 ± 0.0447 [0.1797] | 96 |

Table S2: Segment-level performance of haplotype-specific copy number state identification across simulated ploidy settings. F1 score, precision, and recall are reported for simulated datasets with average ploidy levels of 1.5, 3, 4, and 5. Values are presented as mean ± standard deviation (s.d.), with the median shown in brackets. $n$ denotes the number of successfully evaluated datasets for each method after excluding failed runs. The best-performing method (highest mean value) for each metric within each ploidy level is shown in **bold**.

| Sector | Tool | $n$ | MSE (mean $\pm$ std) |
|---|---|---|---|
| H | Alleloscope | 15 | $0.5514 \pm 0.1672$ |
| | AneuFinder | 24 | $0.3406 \pm 0.0559$ |
| | CHISEL | 24 | $2.7130 \pm 1.7669$ |
| | CNRein | 24 | $1.4715 \pm 0.5584$ |
| | CNVeil | 24 | $\mathbf{0.2563 \pm 0.0194}$ |
| | FLCNA | 24 | $1.0012 \pm 0.1357$ |
| | Ginkgo | 24 | $0.2914 \pm 0.0414$ |
| | HMMcopy | 24 | $3.1281 \pm 0.1376$ |
| | SCOPE | 24 | $0.2919 \pm 0.0510$ |
| | SEACON | 24 | $0.2601 \pm 0.0333$ |
| | SPRINTER | 24 | $0.5307 \pm 0.0683$ |
| | SeCNV | 24 | $0.3164 \pm 0.0558$ |
| A1 | Alleloscope | 18 | $1.1700 \pm 0.1276$ |
| | AneuFinder | 21 | $1.0978 \pm 0.0874$ |
| | CHISEL | 21 | $1.9380 \pm 2.6165$ |
| | CNRein | 21 | $1.5603 \pm 0.5974$ |
| | CNVeil | 21 | $\mathbf{0.8500 \pm 0.0667}$ |
| | FLCNA | 21 | $3.0191 \pm 0.1234$ |
| | Ginkgo | 21 | $1.7501 \pm 0.8189$ |
| | HMMcopy | 21 | $0.9910 \pm 0.0435$ |
| | SCOPE | 21 | $0.8968 \pm 0.0599$ |
| | SEACON | 21 | $1.1244 \pm 0.4826$ |
| | SPRINTER | 21 | $1.8130 \pm 0.0388$ |
| | SeCNV | 21 | $1.0110 \pm 0.0835$ |
| A2 | Alleloscope | 3 | $1.1968 \pm 0.1464$ |
| | AneuFinder | 4 | $0.8441 \pm 0.0683$ |
| | CHISEL | 4 | $1.1761 \pm 0.2020$ |
| | CNRein | 4 | $1.8954 \pm 0.0077$ |
| | CNVeil | 4 | $0.9128 \pm 0.0246$ |
| | FLCNA | 4 | $2.9529 \pm 0.1738$ |
| | Ginkgo | 4 | $1.0554 \pm 0.0691$ |
| | HMMcopy | 4 | $1.1388 \pm 0.0252$ |
| | SCOPE | 4 | $0.7251 \pm 0.0534$ |
| | SEACON | 4 | $\mathbf{0.7047 \pm 0.0677}$ |
| | SPRINTER | 4 | $1.4695 \pm 0.0147$ |
| | SeCNV | 4 | $0.7589 \pm 0.0652$ |
| D | Alleloscope | 42 | $\mathbf{0.0027 \pm 0.0118}$ |
| | AneuFinder | 42 | $0.0569 \pm 0.0116$ |
| | CHISEL | 42 | $3.8685 \pm 0.6839$ |
| | CNRein | 42 | $0.6438 \pm 0.4860$ |
| | CNVeil | 42 | $0.0036 \pm 0.0077$ |
| | FLCNA | 42 | $0.4672 \pm 0.2898$ |
| | Ginkgo | 42 | $0.0450 \pm 0.0163$ |
| | HMMcopy | 42 | $1.0897 \pm 0.1922$ |
| | SCOPE | 42 | $0.0036 \pm 0.0124$ |
| | SEACON | 42 | $0.0104 \pm 0.0125$ |
| | SPRINTER | 42 | $0.0053 \pm 0.0099$ |
| | SeCNV | 42 | $0.0039 \pm 0.0097$ |

Table S3: Values are reported as mean $\pm$ standard deviation (s.d.) for each copy number sector (H, A1, A2, and D) across all evaluated datasets. Lower MSE indicates higher accuracy. $n$ denotes the number of dataset–sector combinations successfully evaluated for each method after excluding failed runs. The best-performing method (lowest mean MSE) within each sector is shown in **bold**.

| Tool | Dataset | CPU Time (Mins) | Memory (GB) |
|---|---|---|---|
| CNVeil | T10 | 71.42 | 81.90 |
| rcCAE | T10 | 1340.93 | 0.38 |
| SeCNV | T10 | 36.82 | 1.43 |
| SCOPE | T10 | 106.73 | 3.26 |
| AneuFinder | T10 | 116.73 | 43.88 |
| HMMcopy | T10 | 32.53 | 3.45 |
| Alleloscope | T10 | 0.18 | 0.10 |
| CNVeil | T16 | 79.50 | 100.00 |
| rcCAE | T16 | 2528.25 | 1.05 |
| SeCNV | T16 | 63.98 | 1.49 |
| CHISEL | T16 | 284.18 | – |
| SCOPE | T16 | 103.45 | 3.50 |
| AneuFinder | T16 | 203.52 | 79.73 |
| HMMcopy | T16 | 35.28 | 3.54 |
| Alleloscope | T16 | 4.70 | 6.99 |
| SEACON | T16 | 1361.28 | – |
| SPRINTER | T16 | 537.75 | 71.59 |
| FLCNA | T16 | 90.15 | 50.00 |
| CNRein | T16 | 26 349.40 | 71.48 |
| CNVeil | KTN302 | 40.35 | 43.82 |
| rcCAE | KTN302 | 1972.92 | 1.84 |
| SeCNV | KTN302 | 35.55 | 0.93 |
| SCOPE | KTN302 | 113.40 | 3.47 |
| AneuFinder | KTN302 | 89.15 | 33.95 |
| HMMcopy | KTN302 | 31.32 | 3.30 |
| Alleloscope | KTN302 | 2.52 | 4.33 |
| CNVeil | sim p1.5 | 10.40 | 0.84 |
| rcCAE | sim p1.5 | 1420.85 | 0.80 |
| SeCNV | sim p1.5 | 12.87 | 1.92 |
| SCOPE | sim p1.5 | 64.15 | 3.88 |
| AneuFinder | sim p1.5 | 58.91 | 40.72 |
| HMMcopy | sim p1.5 | 10.58 | 3.12 |
| CNVeil | sim p3 | 9.95 | 1.23 |
| rcCAE | sim p3 | 4071.77 | 0.62 |
| SeCNV | sim p3 | 13.84 | 1.70 |
| SCOPE | sim p3 | 49.59 | 3.42 |
| AneuFinder | sim p3 | 30.72 | 11.07 |
| HMMcopy | sim p3 | 10.42 | 3.06 |
| CNVeil | sim p4 | 9.99 | 1.47 |
| rcCAE | sim p4 | 1617.10 | 1.13 |
| SeCNV | sim p4 | 12.72 | 1.93 |
| SCOPE | sim p4 | 46.80 | 3.68 |
| AneuFinder | sim p4 | 52.73 | 47.46 |
| HMMcopy | sim p4 | 10.47 | 3.12 |
| CNVeil | sim p5 | 9.57 | 1.31 |
| rcCAE | sim p5 | 4044.58 | 1.39 |
| SeCNV | sim p5 | 13.20 | 1.92 |
| SCOPE | sim p5 | 54.17 | 4.14 |
| AneuFinder | sim p5 | 27.02 | 11.76 |
| HMMcopy | sim p5 | 10.21 | 3.14 |
| CNVeil | TN4 | 91.87 | 98.28 |
| SeCNV | TN4 | 68.02 | 20.77 |
| CHISEL | TN4 | 534.85 | 93.80 |
| AneuFinder | TN4 | 227.23 | 76.75 |
| HMMcopy | TN4 | 32.26 | 3.75 |
| Alleloscope | TN4 | 2.03 | 4.93 |
| SEACON | TN4 | 5400.35 | – |
| Ginkgo | TN4 | 103.95 | 4.69 |
| SPRINTER | TN4 | 1036.72 | 81.79 |
| FLCNA | TN4 | 63.88 | 4.54 |
| CNRein | TN4 | 82 475.45 | 91.61 |

Table S4: **Computing resource consumption of CNV inference tools.** CPU time and peak memory usage are reported for the currently available benchmark results across simulated and real-world datasets. CPU time is reported in minutes, and memory usage is reported in gigabytes. Dashes indicate that peak memory usage could not be recovered from the available Slurm accounting records. For allele- and haplotype-aware methods, the reported CPU time and memory usage begin at the copy number inference step and do not include upstream preprocessing, such as SNP calling and SNP counting.

#### S4 Supplementary Figures

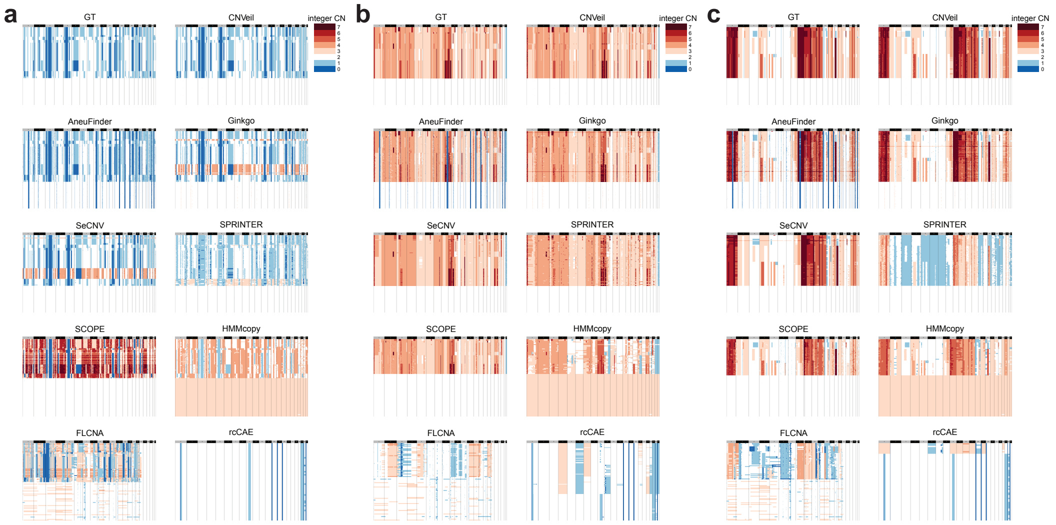

Figure S1: **Representative total copy number inference across simulated datasets with varying ploidy states.** Representative heatmap comparisons of inferred integer copy number profiles across methods for simulated datasets with ploidy states of (a) 1.5, (b) 3, and (c) 5. Columns represent genomic bins ordered by chromosome and rows represent single cells. Ground-truth (GT) copy number states are shown alongside predictions from CNVeil and competing methods.

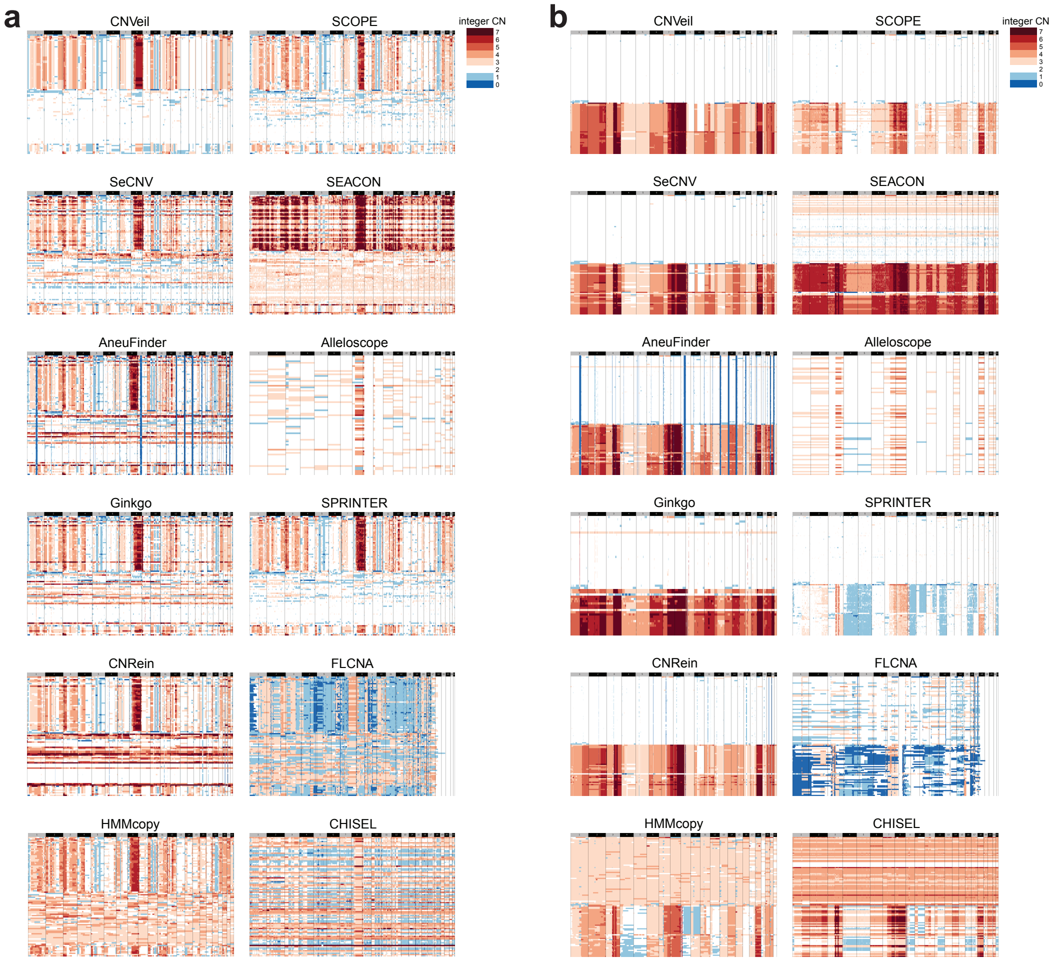

Figure S2: **Extended comparison of total copy number inference across additional single-cell DNA sequencing datasets.** Representative heatmap comparisons of inferred integer copy number profiles across methods for (a) KTN302 and (b) T16. Columns represent genomic bins ordered by chromosome and rows represent single cells. Cells are ordered identically across methods to facilitate direct comparison of chromosome-scale copy number patterns and subclonal organization.

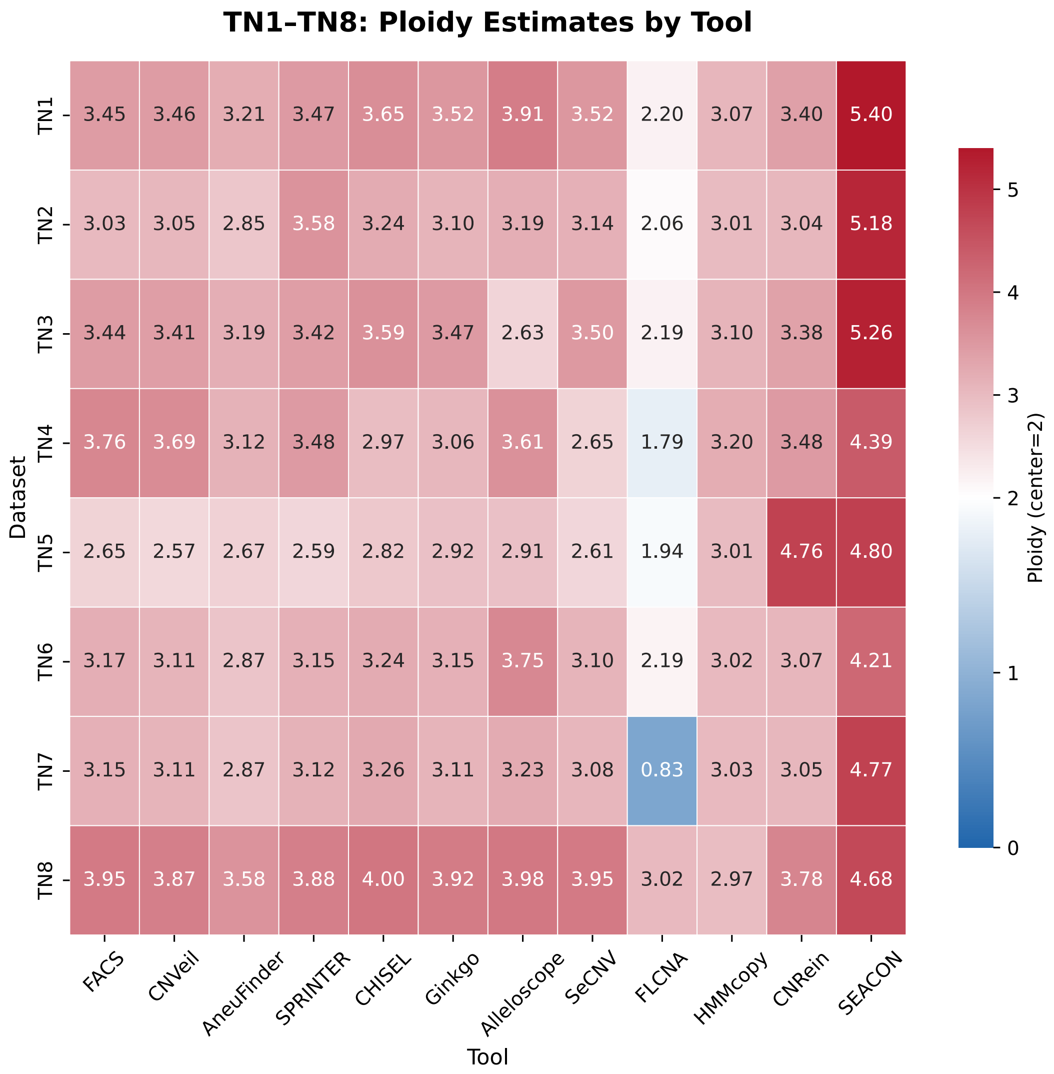

Figure S3: **Comparison of ploidy estimates across ACT samples using FACS and single-cell copy number inference methods.** Heatmap showing genome-wide ploidy estimates across ACT tumor samples (TN1-TN8) inferred by CNVeil and representative single-cell CNV inference methods, together with experimentally measured FACS ploidy values.

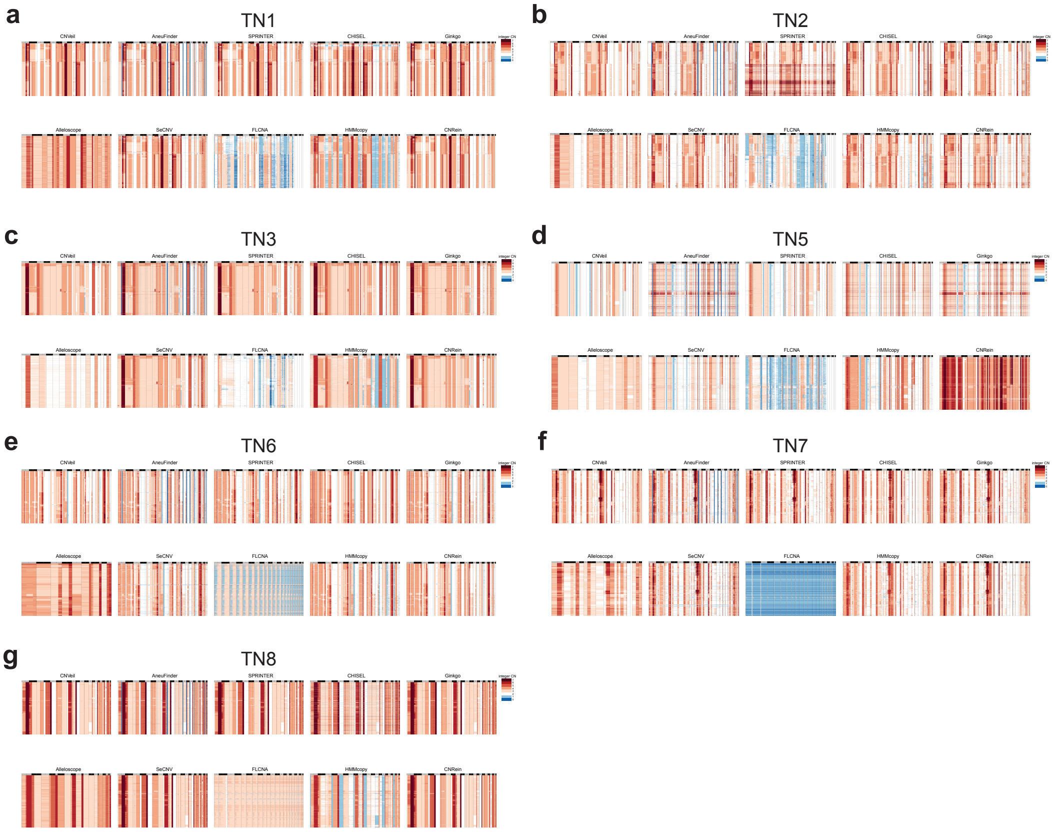

Figure S4: **Extended comparison of total copy number inference across ACT tumor samples.** Representative heatmap comparisons of inferred integer copy number profiles across ACT samples (a-c) TN1-TN3, (d-f) TN5-TN8 for CNVeil and competing single-cell CNV inference methods. Columns represent genomic bins ordered by chromosome and rows represent single cells. Cells are ordered identically across methods to facilitate direct comparison of chromosomal copy number structure and subclonal organization.

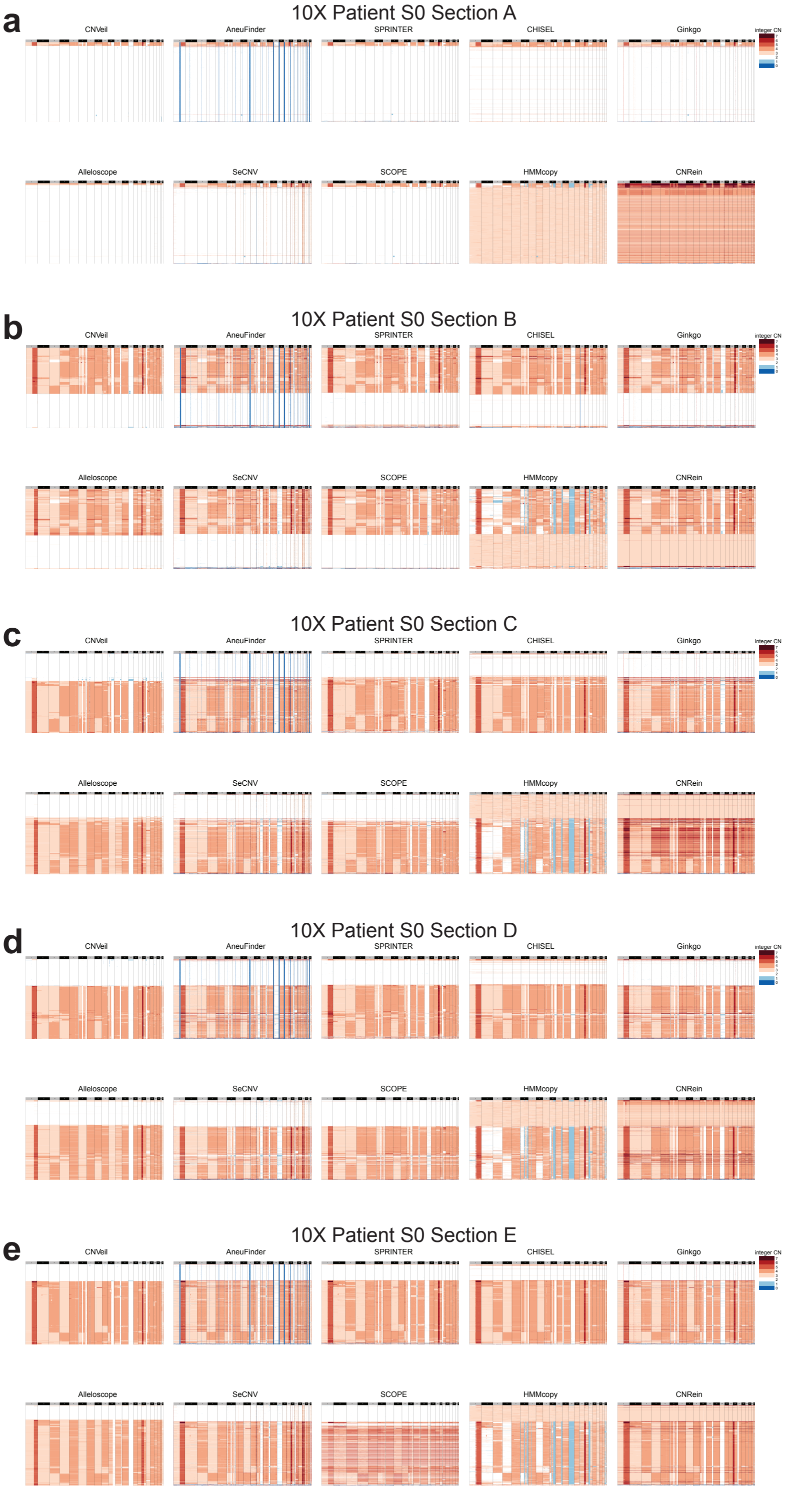

Figure S5: **Extended comparison of total copy number inference across 10x Chromium datasets.** Representative heatmap comparisons of inferred integer copy number profiles across (a-e)10x Chromium Patient S0 Section A-E for CNVeil and competing single-cell CNV inference methods. Columns represent genomic bins ordered by chromosome and rows represent single cells. Cells are ordered identically across methods to facilitate direct comparison of chromosomal copy number structure and subclonal organization.

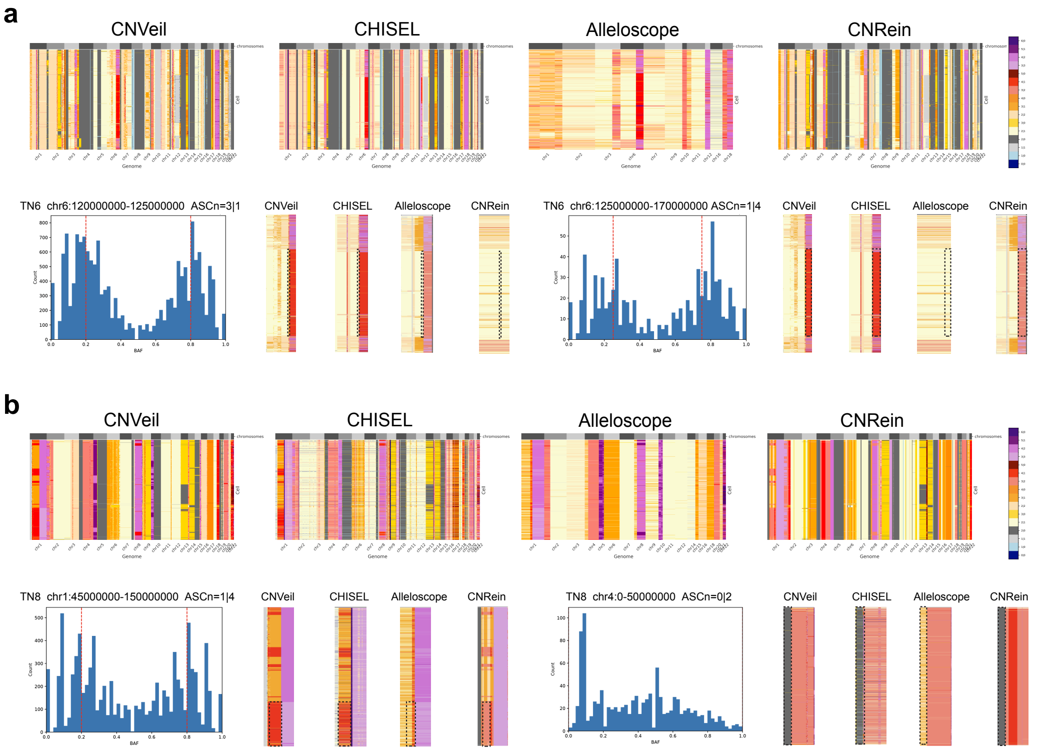

Figure S6: **Extended comparison of allele-specific copy number inference across ACT samples.** (a,b) Representative genome-wide and focal-region comparisons of inferred allele-specific copy number states across methods in ACT (a) TN6 and (b)TN8. Columns represent genomic bins ordered by chromosome and rows represent single cells. Cells are ordered identically across methods to facilitate direct comparison of allelic structure and subclonal organization.

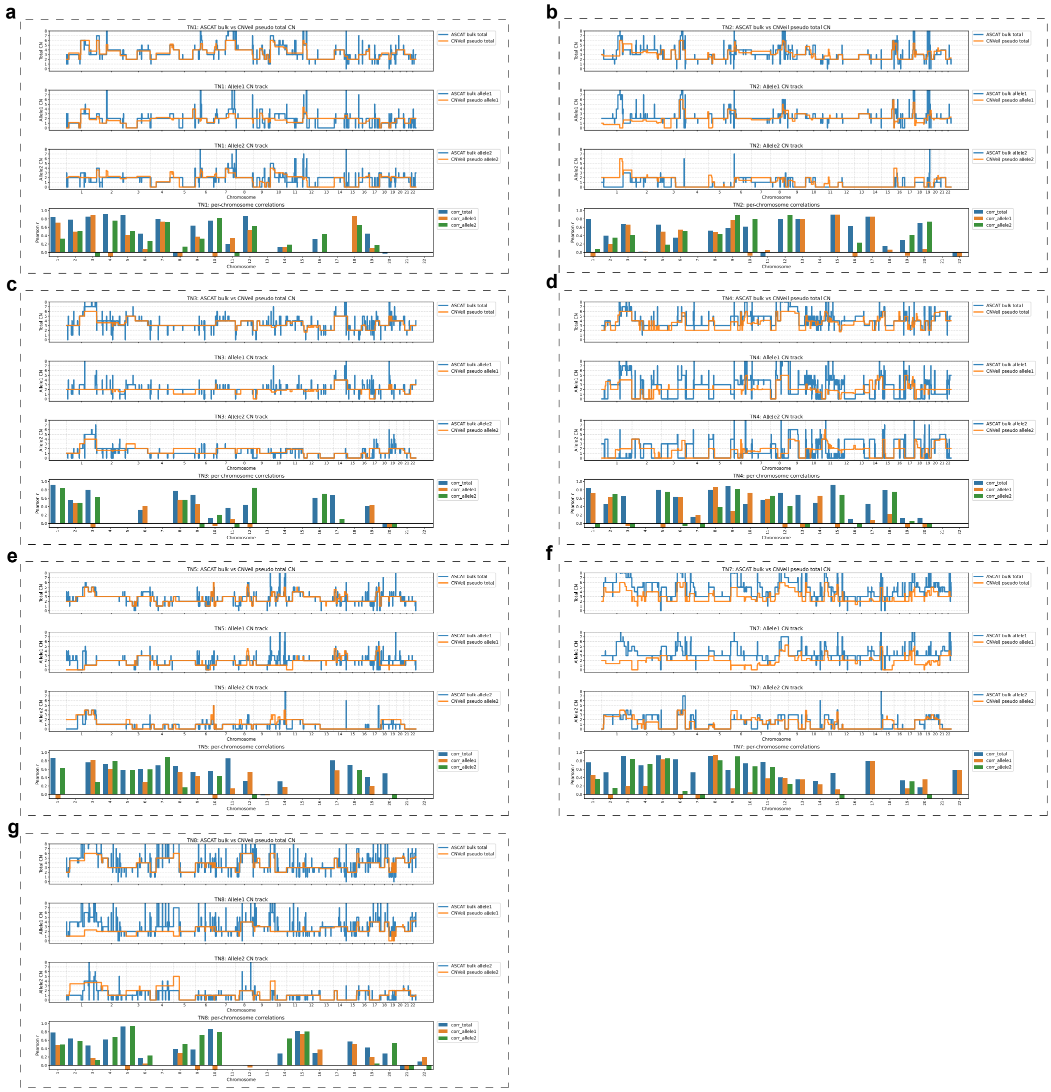

Figure S7: **Genome-wide concordance between CNVeil pseudo-bulk and matched bulk WES haplotype-resolved copy number profiles across ACT samples.** Comparison of pseudo-bulk copy number profiles inferred by CNVeil with matched bulk WES-derived ASCAT profiles across ACT tumors. Panels correspond to individual samples: (a) TN1, (b) TN2, (c) TN3, (d) TN4, (e) TN5, (f) TN7, and (g) TN8. For each sample, the top panel shows total copy number comparison between bulk WES and pseudo-bulk CNVeil profiles, the middle panel shows haplotype-specific copy number comparison, and the bottom panel summarizes chromosome-level Pearson correlations for total and haplotype-specific copy number states.

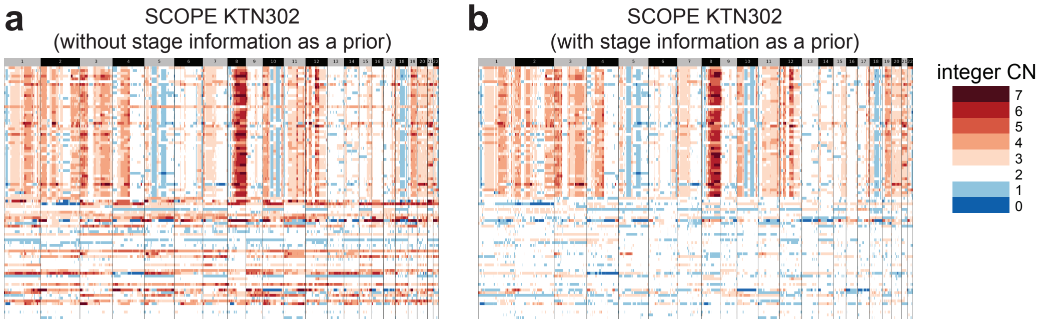

Figure S8: **Effect of prior stage information on SCOPE copy number inference in the KTN302 dataset.** Comparison of SCOPE-inferred integer copy number profiles for the KTN302 dataset (a) without and (b) with treatment-stage information provided as prior knowledge for normal-cell selection. Columns represent genomic bins ordered by chromosome and rows represent single cells. Cells are ordered identically between panels to facilitate direct comparison. Incorporating stage information improves identification of the diploid mid-treatment population and reduces the number of incorrectly inferred hyperdiploid cells.
